# Proximity interactomics identifies RAI14, EPHA2 and PHACTR4 as essential components of Wnt/planar cell polarity pathway in vertebrates

**DOI:** 10.1101/2025.01.15.633117

**Authors:** Kristína Gömöryová, Nikodém Zezula, Tomasz W. Radaszkiewicz, Petra Paclíková, Štěpán Čada, Kateřina Hanáková, Ranjani Sri Ganji, Miroslav Micka, David Potěšil, Zbyněk Zdráhal, Vítězslav Bryja

## Abstract

Wnt/planar cell polarity (Wnt/PCP) pathway is an evolutionarily conserved signaling cascade playing an inevitable role in cell biology. Deregulation of Wnt/PCP leads to severe developmental defects or cancer progression. Here, we applied proximity-dependent biotinylation (BioID) to capture the intracellular interactome of key Wnt/PCP components: transmembrane ROR1, ROR2 and VANGL2, and cytoplasmic DVL3 and PRICKLE1. Mapping of individual preys across the baits and subcellular compartments identified a group of 30 proteins that we tested by loss-of-function in zebrafish. Among those rai14, epha2 and phactr4 were essential for several Wnt/PCP-dependent processes such as zebrafish convergent extension, orientation of lateral organ cells, or migration of melanoma cells. Mechanistically, RAI14, EPHA2 and PHACTR4 connect the receptor complex to effector actomyosin. In summary, this study identified novel essential components of vertebrate WNT/PCP pathway and provides a detailed characterization of PCP complex composition that can serve as comprehensive resource for further analysis of Wnt/PCP function.

## INTRODUCTION

Establishment and maintenance of cell polarity is critical for the correct development of all multicellular organisms. In addition to apicobasal polarity, cells are aligned also orthogonally, within the plane of the tissue via a mechanism called planar cell polarity (PCP)^1^. PCP genes were initially identified in the fruit fly *Drosophila melanogaster* genetic screens, but currently the PCP machinery is recognized as an evolutionary conserved module that controls cell polarization in all animals^2,3^. The ‘core’ PCP system is centered around Wnt-Frizzled signaling (Wnt/PCP signaling)^4^. Wnt/PCP pathway is the best described, evolutionarily and functionally conserved, branch of the non-canonical Wnt signaling^4^. In contrast to the canonical Wnt (Wnt/β-catenin) signaling, stimulation of the Wnt/PCP pathway leads to the activation of small GTPases and subsequent cytoskeleton reorganization or cell migration rather than regulation of gene transcription^1,5,6^. In the fruit fly wing epithelium, the polarity is established by asymmetrical localization of Wnt/PCP proteins Van Gogh (Vang) and Prickle (Pk) at the proximal side of the cell, and Frizzled (Fz), and Dishevelled (Dsh) at the distal side of the cell. The PCP pattern is propagated across the tissue and stabilized by homophilic interactions of the protein Flamingo, located both proximally and distally^1,7^.

Unlike in *Drosophila*, in vertebrates several homologues of the core PCP proteins are present: e.g. there are ten Frizzled (Fzd1-10), four Prickle (Prickle1-4), three Dishevelled (Dvl1-3) and two Vang-like (Vangl1/2) proteins^1,3,4,8,9^. Despite the fact the core PCP module composed of the PCP proteins mentioned above seem to be conserved, there are several other proteins associated with the Wnt/PCP signaling in vertebrates. These include especially dedicated Wnt ligands WNT5A and WNT11, and atypical receptor tyrosine kinases ROR1, ROR2, RYK and PTK7 that were proposed to act as their (co-)receptors^8,10–12^.

Mutations or other alterations of Wnt/PCP genes in human and other vertebrates lead to a variety of developmental defects. Many of them fall into a category of neural tube closure defects, such as spina bifida, exencephaly or craniorachischisis^13,14^ and reflect the absolute requirement of Wnt/PCP pathway during the process of convergent extension (CE) that precedes neural tube closure^1,4^. Other type of defects, such as Robinow syndrome, a congenital skeletal dysplasia^15^ demonstrates the key role of Wnt/PCP in later morphogenetic events during organ development^1^. Additionally, given the major role of cell polarization coordinated by Wnt/PCP in cell migration, it is of no surprise that dysregulation of the Wnt/PCP has been linked to a variety of aggressive cancers, where Wnt/PCP pathway triggers cell invasion, treatment resistance and metastasis^16^.

The molecular details of Wnt/PCP in mammalian cells are largely unknown. In order to address this gap, we have employed the proximity-dependent biotinylation (BioID)^17^ to extensively characterize the interactome of selected Wnt/PCP proteins in T-REX 293 cells. The unbiased interactomics of transmembrane ROR1, ROR2, VANGL2, and cytoplasmic DVL3 and PRICKLE1 allowed us to describe general features of core PCP module in human cells. Functional screening of the interacting proteins in zebrafish identified three proteins – rai14, epha2 and phactr4 – as essential regulators of PCP *in vivo*. Follow-up mechanistic studies showed that the function of EPHA2, PHACTR4, and RAI14 is conserved in WNT5A-dependent migration of human melanoma cells where these proteins appear essential for the functional connection of the receptor complex and the actomyosin cytoskeleton.

## RESULTS

### Proximity interactomes of ROR1, ROR2, VANGL2, PRICKLE1 and DVL3

In order to characterize the interactome of the selected human Wnt/PCP pathway components – Wnt co-receptors ROR1, ROR2 and transmembrane polarity regulator VANGL2, as well as cytoplasmic proteins DVL3 and PRICKLE1 – we employed the proximity-dependent biotinylation (BioID)^17^. BioID takes advantage of the promiscuous biotin ligase BirA*, which is able to biotinylate proteins in a radius of approximately 10 nm around the protein of interest (POI, bait). BioID thus allows to capture both stable and transient protein interactors (preys). We prepared T-REx 293 TetON cell lines that inducibly express the POI fused with BirA* ligase and tag (either HA or MYC, Fig. 1A). The biotinylation in these cells was specific, i.e. POI-BirA* biotinylated proteins in proximity only in the presence of tetracycline (Tet) and biotin (Suppl. Fig. 1A-E). For each cell line/POI we performed biotinylation and subsequent streptavidin pulldown in 6 biological replicates. All samples were then subjected to mass spectrometry (MS) analysis. To generate the high-stringency protein interaction list, we filtered out the proteins that non-specifically bound to BirA* (using the CRAPOME database)^18^ and proteins not detected in at least 4 replicates of the same bait. This approach resulted in a list of 985 high-confidence preys across all baits (Suppl. Table 1).

**Figure 1.**
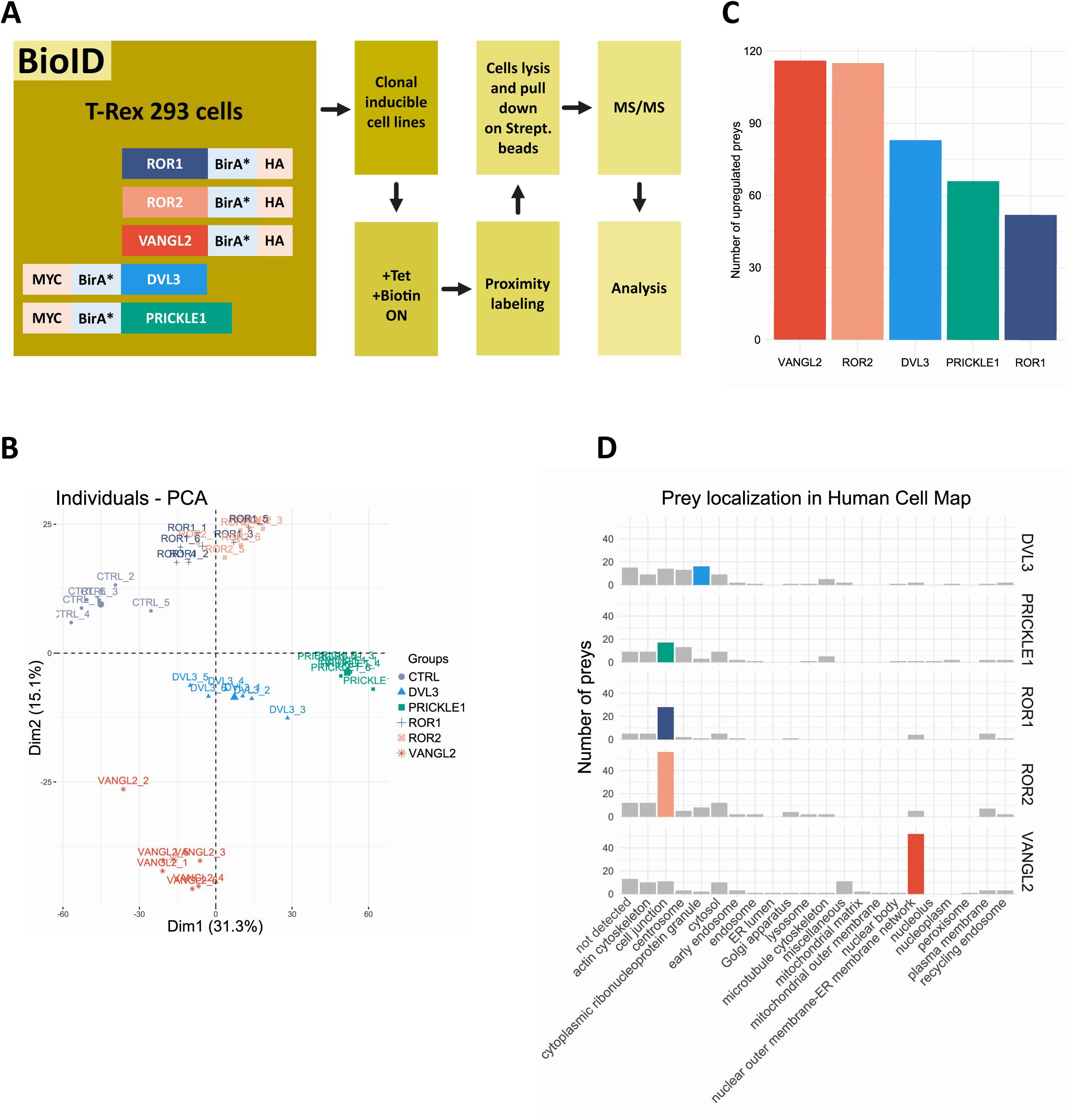
Proximity interactomes of ROR1, ROR2, VANGL2, PRICKLE1 and DVL3. **A.** Schemes of constructs representing the core Wnt/PCP components and overview of the experiment workflow Human proteins were tagged with BirA*, and MYC (DVL3, PRICKLE1) or HA (ROR1, ROR2, VANGL2), respectively. T-REx 293 cell line was stably transfected with selected and inducible constructs, and upon addition of biotin, interactors within the proximity of bait were labelled. After streptavidin pulldown, samples were subjected to LC-MS/MS. Following data analysis revealed several new Wnt components interactors, which PCP phenotype were further tested both in cell lines and zebrafish. **B.** Principal Component Analysis showing distinct clusters corresponding to particular baits, whereas individual replicates of particular baits cluster together. ROR1 and ROR2 samples highly overlap, suggesting overlap also in their interactomes. **C.** Number of upregulated preys (denoted based on log_2_ fold change >1 and adjusted p-value <0.05) for particular baits. **D.** Cellular localization of upregulated preys based on the Human Cell Map resource shows distinct localization patterns of baits interactors. ROR1 and ROR2 interactions occur predominantly in the cell junctions, DVL3 interactions in the cytoplasmic ribonucleoprotein granules, PRICKLE1 interactions in the cell junctions and centrosome, and VANGL2 interactions in the endoplasmic reticulum.

Principal component analysis of remaining protein intensities in individual replicates (PCA, Fig. 1B) excluded the batch effect in the data and at the same time suggested the existence of biological differences among baits with the exception of ROR1 and ROR2 that had similar interactomes. Differential expression analysis (based on cut off: adj. p value < 0.05 and log_2_ fold change >1 or <-1) identified the individual preys for each bait (Suppl. Fig. 1F, Suppl. Table 2). VANGL2 and ROR2 had the highest number of interactors (∼100), whereas ROR1 reported the lowest number of interacting proteins (∼50) (Fig. 1C). Visualization by UpSet plot (Suppl. Fig. 1G, Suppl. Table 3) showed that there are only 18 interactors shared among all 5 baits, including DVL2 and DVL3. On the contrary, VANGL2 has the highest number of unique interactors (69), followed by ROR2 (42). Gene ontology (GO)^19^ analysis (Suppl. Fig. 1H) showed that interactome of all baits is enriched in proteins associated with the *Wnt canonical pathway*, *cell division* and *cytoskeleton organization*, including *RHO GTPase cycle*. The most significant bait-specific features likely associate with the unique biosynthesis of VANGL2 (GO terms initiation of: *nuclear envelope reformation* and *insertion of tail-anchored proteins to endoplasmic reticulum membrane*) and capacity of DVL3 to form protein condensates (GO term: *stress granule assembly*). Using the BioID-based Human Cell Map (HCM)^20^ that is built on the same cell line, we could also map individual interactors into the cellular compartments (Fig. 1D). ROR1 and ROR2 preys associated predominantly with the cell junction and plasma membrane, whereas DVL3 interactors most commonly localized into cytoplasmic ribonucleoprotein granules. In case of PRICKLE1, it was primarily the cell junction and centrosome. Most VANGL2 preys localized to the nuclear outer membrane/endoplasmic reticulum network, which likely does not reflect the interactors mediating the action of mature VANGL2, but rather represent proteins that are associated with VANGL2 biosynthesis and maturation.

### Global features of the interactome of core mammalian PCP proteins

We hypothesized that possible novel regulators of PCP shall interact with two or more baits. To explore the interactome in this direction, we performed *k*-means clustering of all upregulated proteins considered as prey of at least one bait protein (Fig. 2A, Suppl. Table 4). This analysis revealed that there are roughly five patterns of bait-prey interaction, visible as protein clusters: cluster (#C) 1, composed mostly of DVL3 and PRICKLE1 interactors; #C2 composed of almost exclusively VANGL2 interactors; #C3 composed of ROR1 and ROR2 interactors; #C4 composed of ROR1, ROR2 and PRICKLE1 interactors, and #C5 composed of interactors common for all baits.

**Figure 2.**
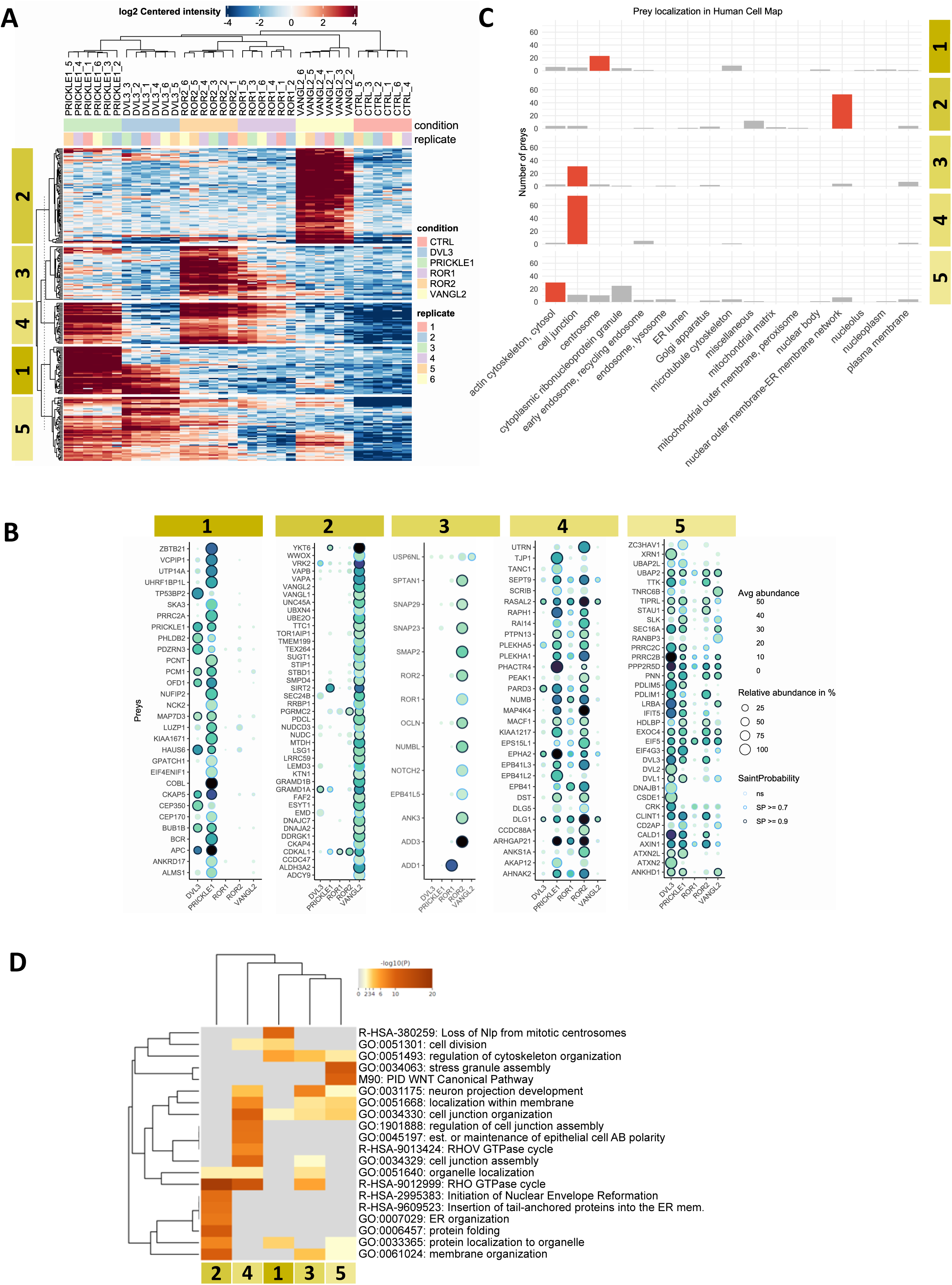
Global features of the interactome of core mammalian PCP proteins. **A.** Heatmap of proteins upregulated in at least one bait. Approximately 5 different clusters of preys are detected using k-means clustering. **B.** Dotplots displaying the results of SaintExpress analysis, verifying the bait-prey interactions for individual clusters. Dot color corresponds to abundance (the darker color, the more abundant was prey), size - to the relative abundance, edge color represents the SaintProbability (the darker color, the stronger was bait-prey interaction). **C.** Human Cell Map analysis of preys localization in the individual clusters. Cell localization with the most preys is labeled in red. **D.** Gene ontology analysis for the individual clusters.

As a complementary scoring approach that considers the number of MS/MS spectra rather than intensity, we used the SAINTexpress^21^ analysis within the REPRINT^18^ resource (Suppl. Table 5). We visualized the results in a dotplot for each cluster using strict filtering criteria with the SAINT probability (SP) of interaction higher than 0.9 (Fig. 2B) This analysis confirmed the cluster composition, where preys identified by clustering method in a previous step predominantly interact with estimated baits. The complete dotplot with all interactions that passed the threshold, as well as another, showing non-significant interactions, can be found in the Supplementary information (Suppl. Fig. 2A-B).

Human Cell Map analysis showed striking differences among the clusters. #C1 (DVL3 and PRICKLE1) preys localize predominantly to the centrosome, VANGL2 interactors from #C2 occur in the nuclear outer-membrane ER network, interactors of ROR1, ROR2 and PRICKLE1 (#C3 and #C4) localize primarily to the cell junction, and interactors of shared baits (#C5) are distributed all across the compartments, with the highest number of preys localizing to the actin cytoskeleton and cytosol (Fig. 2C). This subcellular localization of preys in individual clusters is reflected also in the GO analysis (Fig. 2D).

### Functional testing in zebrafish identifies RAI14, PHACTR4 and EPHA2 as essential components of PCP machinery

Based on the subcellular localization as well as GO ontology, we decided to functionally focus on the proteins from #C4. Proteins in this cluster interact primarily with PCP-dedicated baits ROR1, ROR2 and PRICKLE1, are predominantly localized at the cell junction, where PCP signaling takes place and are enriched in GO terms previously associated with PCP such as *establishment of epithelial polarity* or *RHO GTPase cycle*. In order to test the function of individual #C4 proteins we have performed a loss-of-function screen in zebrafish, *Danio rerio*. Zebrafish is a suitable model to study Wnt/PCP whose disruption results in a variety of developmental defects, such as defective sensory hair cell orientation of lateral line organ, shortened and widened body axis and neural tube closure defect^1,22,23^. To test the role of a protein in the WNT/PCP pathway, we utilized the CRISPR/Cas9 approach to produce in zebrafish F0 generation knockouts (KOs), termed “crispants”, of the genes encoding #C4 proteins and analyzed the presence of defects associated with the disrupted PCP (Fig. 3A).

**Figure 3.**
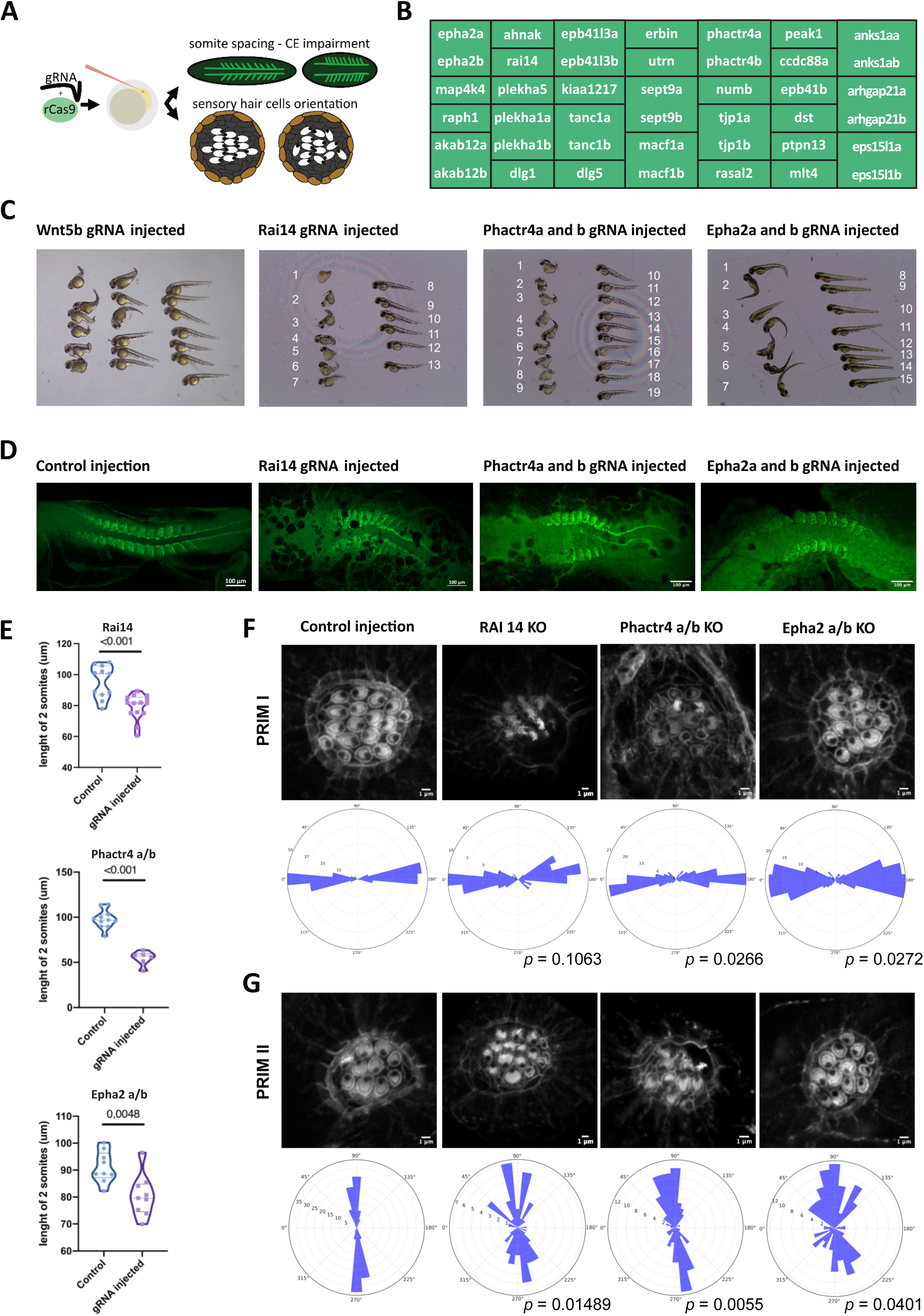
Planar Cell Polarity *in vivo* - functional screening. **A.** Simplified scheme of *in vivo* testing with representation of typical phenotypes of impaired WNT/PCP pathway. **B.** List of tested zebrafish genes from cluster 4. **C.** Overall phenotype of *rai14*, *phactr4a/b* and *epha2a/b* crispants at 2 dpf. **D.** Representative images of myoD1 whole mount immunofluorescence, labeling somitogenesis in control, *rai14* KO, *phactr4a/b* KO and *epha2a/b* KO 14.5 hpf zebrafish embryos. **E.** Analysis of convergent extension impairment. The length of the first 2 anterior somites was measured for control embryos and *rai14*, *phactr4a/b* and *epha2a/b* KO embryos. **F.** Representative images and rose plots of the degree of polarization of sensory hair cells of PRIM I and PRIM II derived neuromasts in control, *rai14*, *phactr4a/b* and *epha2a/b* KO embryos. The level of significance was measured using axial representation of the data and tested with Levene’s test. n values: PRIM I (Control n=6, epha2 n=6, phactr4 n=5, rai14 n=5) PRIM II (Control n=7, epha2 n=7, phactr4 n=6, rai14 n=4).

Out of the 33 proteins from #C4 we functionally tested 30. The three excluded genes are *SCRIB* as it has been already shown to regulate PCP^1^, *PARD3* because it had more than two homologues present in the fish and *AHNAK2* as it had none. Remaining 30 human genes corresponded to 42 zebrafish orthologues (**Fig. 3B**). In cases where two homologues of the same gene existed, both were targeted simultaneously. First, we have optimized the screening system using KO of *wnt5b*, which has in zebrafish a well-defined PCP phenotype. As shown in **Fig. 3C**, the *wnt5b* crispants show clearly shortened body axis in 2-day post-fertilization (2dpf) embryos in approximately 50% of cases. This incomplete penetrance is probably due to the combination of mosaicism and non-productive targeting. Embryos injected with a control solution show similar phenotypes in max. 5% of cases. Still, this assay provided sufficient readout and we considered a hit the situation where PCP phenotype was observed in 25-50% of injected embryos. Based on these criteria we were able to functionally test zebrafish orthologous genes using the gRNAs summarized in Suppl. Table 12. In 2dpf embryos we observed shortened body axes in *rai14*, *phactr4a/b* and *epha2a/b* KOs (Fig. 3C). Similar phenotype has been observed also in some biological replicates for *tanc1a/b*, however, this result was inconsistent and we thus excluded *tanc1a/b* crispants from the follow up analysis.

To further confirm that the phenotypes observed in the KO of *rai14*, *phactr4a/b*, and *epha2a/b* are indeed caused by the defects in the convergent extension (CE), we examined the process of somitogenesis in 14-hour post-fertilization (hpf) old embryos (Fig. 3D, Suppl. Fig. 3A). Disruption of *rai14*, *phactr4a/b*, and *epha2a/b* resulted in the shorter somites, which is a typical hallmark of the impaired CE (Fig. 3E). To provide additional proof of the essential role of these proteins in the WNT/PCP signaling pathway, we investigated the PCP-dependent polarity of the sensory hair cells in the lateral line organ. We have focused on two types of neuromasts present in the lateral line organ. In type PRIM I, that is derived from primordium I, the longitudinal alignment of sensory cells depends on core PCP proteins. In type PRIM II, that develops later from primordium II, the dorso-ventral orientation of sensory cells oriented, depends more on the activity of the Wnt/receptor complex^22,24^. As shown in Fig. 3F, G and Suppl. Fig. 3B, the polarity of the sensory hair cells in PRIM II has been disrupted by the KO of all three genes *rai14*, *phactr4a/b*, and *epha2a/b*. Only *epha2a/b* KO affected also the polarity in PRIM I. Of note, in *rai14* crispants, also the overall morphology was affected with a very small number of sensory cells within each organ (Suppl. Fig. 3B, Fig. 3F), indicating a more general role of *rai14* in the development of the lateral line organ. Nevertheless, using two independent readouts, our data support essential function of RAI14, PHACTR4 and EPHA2 in the vertebrate Wnt/PCP pathway. Strong phenotypes in the Wnt-dependent PRIM II polarity^22^ suggest that they functionally act between the receptor complex and downstream effectors.

### The interactome of RAI14, PHACTR4 and EPHA2 overlaps with the interactome of core PCP proteins

Experiments conducted in the zebrafish model allowed us to describe RAI14, PHACTR4 and EPHA2 as novel regulators of WNT/PCP. These three proteins have very different domain compositions (**Fig. 4A**). RAI14 has been characterized for the first time in 2001 by two groups of researchers^25,26^. One of the names of this protein – Ankycorbin (Ankyrin repeat- and coiled-coil structure-containing protein) refers to the seven ankyrin repeats residing within its N-terminal part and C-terminal coiled-coil folds. These domains, together with the N-terminal amphipathic helix, mediate binding to the plasma membrane and its subsequent shaping; a process shown to be crucial for neuromorphogenesis^27,28^. Its other name – RAI14 (Retinoic acid induced 14) refers to the fact that retinoic acid treatment upregulates its expression in human retinal pigment epithelial cells^26^. Phosphatase And Actin Regulator 4 (PHACTR4) contains disordered regions and three RPEL (RPxxxEL) repeats. This domain is required for the non-polymerized actin biding, suggesting potential involvement of PHACTR4 in the regulation of the cell shape and cytoskeleton^29^. Ephrin type-A receptor 2 (EPHA2) is a receptor tyrosine kinase (RTK) which acts as a receptor in the ephrin signaling pathway. EPHA2 acts also noncanonically via ligand independent manner, involving dimerization with other signaling molecules, like EGFR or E-Cadherin^30,31^. EPHA2 has a single transmembrane (TM) domain and extracellular N-terminus with ligand-binding domain (LBD), Sushi domain, an epidermal growth factor-like motif within a cysteine-rich domain (CRD) and two fibronectin-type III repeats (FN III1 and FN III2). Intracellular sequence contains a juxtamembrane region (JM), kinase domain (KD), sterile α motif (SAM) and a PDZ binding sequence.

**Figure 4.**
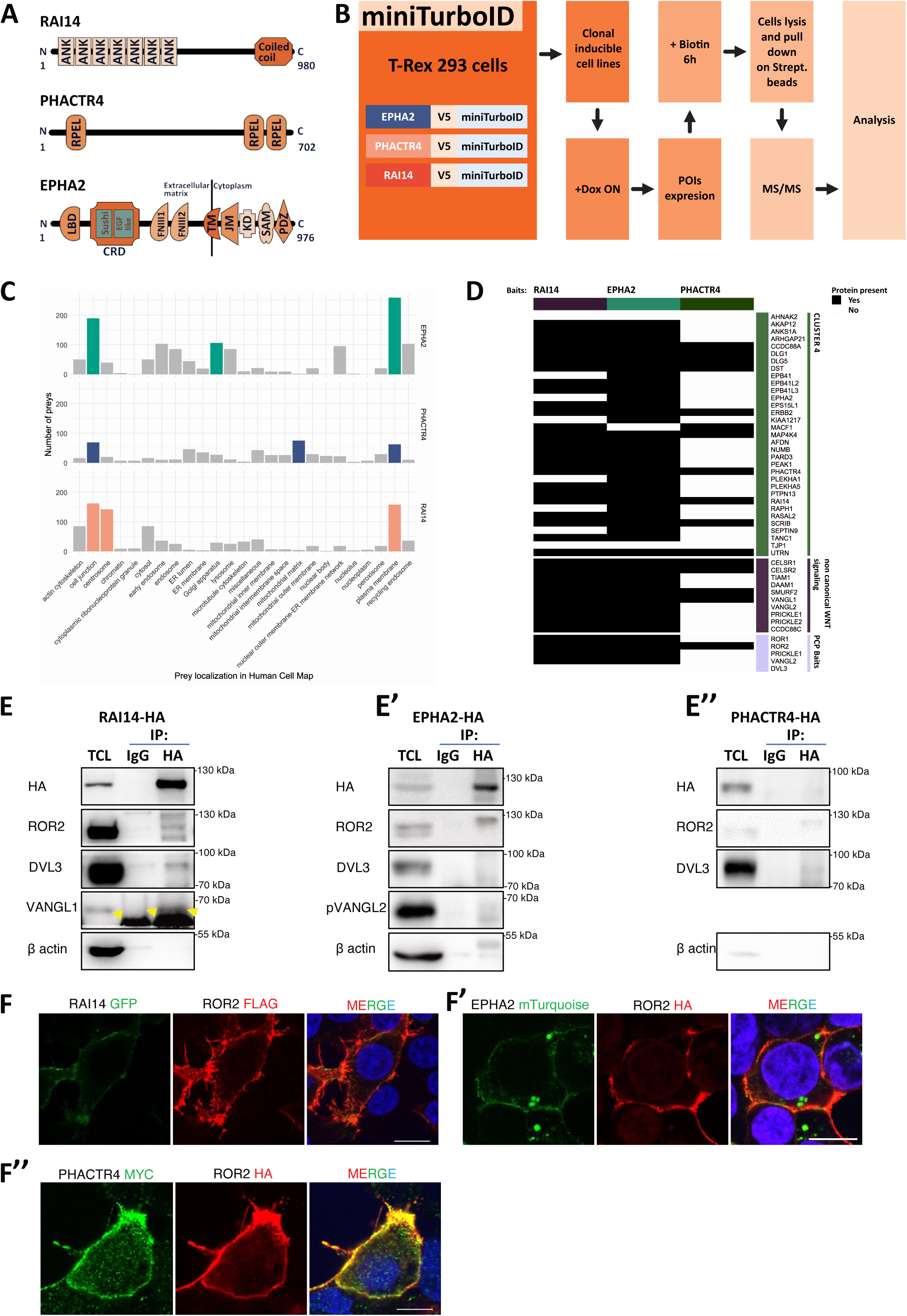
The interactome of RAI14, PHACTR4 and EPHA2 overlaps with the interactome of core PCP proteins. **A.** Schematic representation of the domain structure of the three selected candidate proteins: RAI14, EPHA2 and PHACTR4. **B.** Simplified overview of the miniTurboID experiment. **C.** Human Cell Map analysis of the upregulated preys for three baits, most prominent localization is labeled. All baits show strong localization pattern in the plasma membrane and cell junction, respectively. **D.** Heatmap depicting interaction of 3 baits with the (i) original BioID baits; (ii) proteins from the cluster 4; (iii) non-canonical Wnt components. EPHA2 and RAI14 exhibit similar binding pattern, whereas PHACTR4 interacts only with a subset of proteins. **E.** Immunoprecipitation of HA-tagged RAI14 (E), EPHA2 (E’), PHACTR4 (E’’). Expression of fusion proteins in T-REx 293 stable cell lines was induced by ON treatment with doxycycline. Pull-down using anti-HA tag antibody and IgG control were performed to validate interaction of POIs with components of the Wnt/PCP pathway, N=3. Arrowhead (E) points VANGL1 specific signal. **F.** T-REx 293 cells transiently overexpressing ROR2 and RAI14-GFP (F), EPHA2-mTurquoise (F’) and PHACTR4-MYC (F’’). Staining and confocal microscopy showed colocalization of POIs with ROR2 co-receptor. Scale bars represent 10 μm.

In order to get better mechanistic insight into their function, we have returned back to the mammalian cell culture. Since not much is known about the biology of these proteins, we performed another unbiased interactomics screen using the miniTurboID method (Fig. 4B). MiniTurboID^32^ is based on the very same principle as BioID, but instead of BirA*, it employs a smaller and more efficient miniTurboID ligase. We prepared T-REx 293 TetON cell lines inducibly expressing EPHA2, PHACTR4 and RAI14 tagged with V5 and miniTurboID at their C-termini. This approach allowed us to analyze not only interactomes of these proteins but also describe their subcellular localization. RAI14 and EPHA2 displayed mainly plasma membrane localization, while PHACTR4 was detected in the cellular junctions and cytoplasm (Suppl. Fig. 4A). As shown in Suppl. Fig. 4A and B biotinylation was detected only upon biotin supplementation and transgene expression activation. TurboID followed by LC-MS/MS was then performed in five biological replicates.

More effective biotinylating enzyme as well as more advanced acquisition method (in comparison to original baits in Fig. 1), resulted in a detection of a larger number of biotinylated proteins (Suppl. Table 8) and a higher number of confirmed preys (based on the threshold of adjusted p-value <0.05 and logFC >1) for individual baits (Suppl. Fig. 4D, Suppl. Table 9). Most of the preys have not been previously reported in the Human Cell Map (HCM) probably because HCM is based on less advanced version of the BioID assay. However, the analysis of the preys reported in HCM (Fig. 4C) showed that for all RAI14, EPHA2 and PHACTR4 interactors the plasma membrane and cell junction were among the two out of three most abundant compartments. This is in concordance with the GO analysis (Suppl. Fig. 4E) as well as with the previous observation (Fig. 2C), in which most of the #C4 proteins localized to the cell junction. Importantly, RAI14, EPHA2 and PHACTR4 interacted with (i) original baits used in the BioID, (ii) other conserved components of the non-canonical Wnt pathway and (iii) #C4 proteins (Fig. 4D). Overall, RAI14 and EPHA2 interactome contained more PCP-relevant and #C4 proteins than the interactome of PHACTR4. RAI14 and EPHA2 interacted with all 5 original baits, other conserved PCP proteins and, out of 33 proteins in the cluster #C4, EPHA2 interacted with 30 and RAI14 with 25. PHACTR4 was found to interact only with ROR2, CELSR1/2, SMURF2 and VANGL1 and with 11 proteins in the #C4 (Suppl. Table 10).

In summary, the proximity interactomics confirmed that RAI14, PHACTR4 and EPHA2 interact with multiple known components of PCP pathway as well as with the putative novel regulators from cluster #C4. In order to validate the interactions of PCP proteins with RAI14, PHACTR4 and EPHA2 using independent method, we have produced T-REx 293 cells with doxycycline-inducible expression of HA-tagged variants of these three POIs. These cells have been then used for co-immunoprecipitation experiment using HA pulldown. In case of RAI14, we could confirm a stable interaction with endogenous ROR2, DVL3, and VANGL1 (Fig. 4E). Interestingly, DVL3 has not been detected in the miniTurboID experiment. The presence of this protein in the pull-down sample might thus be a result of the ROR2-DVL3 secondary interaction. EPHA2-HA formed complexes with ROR2 and phosphorylated VANGL2 (Fig. 4E’), whereas PHACTR4 interacted with ROR2 and DVL3 (Fig. 4E’’). Interactions of RAI14, EPHA2 and PHACTR4 to ROR2 were additionally confirmed by a co-localization assay (Fig. 4F-F’’). These data validate the BioID findings and establish that ROR2 co-receptor is a common interactor of RAI14, EPHA2 and PHACTR4.

### RAI14, EPHA2 and PHACTR4 and are required for Wnt5a-dependent cell migration and link membrane complexes to effector actomyosin

To address the role of novel PCP regulators in mammalian cells, we selected melanoma cell line A375 as a well-established model where non-canonical Wnt/PCP module triggered by WNT5A/ROR2 complex controls cell migration and invasion^33^. First, we generated A375 lines lacking RAI14, EPHA2 and PHACTR4 proteins using the CRISPR/Cas9 technology (Fig. 5A). Validation of KO cell lines was performed by NGS (Suppl. Table 7), and in case of RAI14 and EPHA2 by Western blot (Fig. 5A and 5A’). Unfortunately, we were unable to obtain any anti-PHACTR4 antibody that would specifically detect endogenous PHACTR4. All KO cells were able to respond to WNT5A – visualized as DVL2 phosphorylation shift in SDS-PAGE – similarly to parental wild type (WT) A375 line (Fig. 5A-5A’’). This suggests that neither RAI14, EPHA2 nor PHACTR4 are required for the functional activation of the receptor complexes.

**Figure 5.**
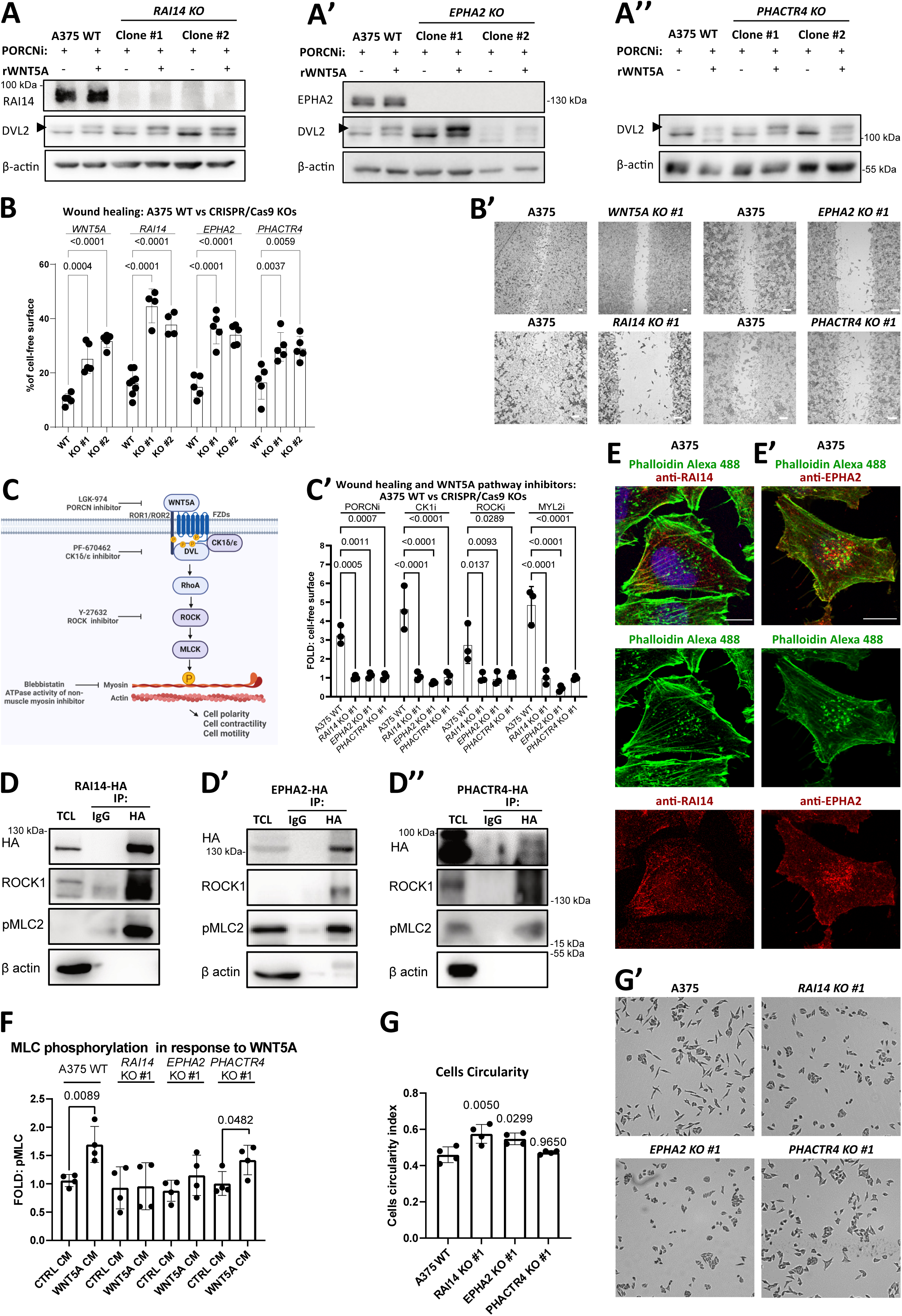
RAI14, PHACTR4 and EPHA2 are required for Wnt5a-dependent cell migration. **A.** Analysis of responses to recombinant WNT5A treatments of A375 wild type cells in comparison to *RAI14* KO (A), *EPHA2* KO (A’) and *PHACTR4* KO (A’’) CRISPR/Cas9 derivate. Two independent clones were used for each protein of interest. Overnight LGK-974 (PORCN1i) 1 µM was used to block the endogenous Wnt secretions and 3 hours long incubation with 100 ng/ml recombinant WNT5A to trigger cellular responses. Arrowheads mark phosphorylation-depended electrophoretic shift of DVL2. Actin β is a loading control, N=3. **B.** Wound healing assay performed on A375 parental and *WNT5A*, *RAI14* and *EPHA2* KO cells. Overnight serum starved cells were wounded. After 48 h cells were fixed, stained and analyzed. Statistical assessment was performed by one-way Anova test, N>=5, *p* values are shown. Representative photos are presented (B’), scale bars = 100 µm. **C.** Schematic representation of the Wnt5a signaling. Binding of the Wnt5a ligand to Frizzled receptors (FZD) and ROR1/2 co-receptors triggers cytosolic cascade of events. Casein kinase 1δ/ε (CK1δ/ε) activates Dishevelled (DVL) and receptor complexes. This results in the signal propagation to the Rho/ROCK, resulting in cytoskeletal rearrangements. This axis can be targeted by i.e. prevention of Wnt ligands secretion (Porcupine inhibition by i.e. LGK-974), enzymatic activity inhibition of CK1δ/ε (PF-670462), ROCK (Y-27632) or myosin (Blebbistatin). Scheme created by Biorender. These interventions had no significant impact on the *RAI14*, *EPHA2* and *PHACTR4* KO A375 cells motility in the wound healing assay in comparison to the parental control (C’). Overnight starved cells were wounded, treated with 1 µM LGK-974, 3 µM PF-670462, 10μM Y-27632 or 5μM Blebbistatin. Cells were fixed, stained and analyzed 48 h after scratch, one-way Anova test, N=3, *p* values are indicated. Graph represents inhibition fold. **D.** Immunoprecipitation of HA-tagged RAI14 (D), EPHA2 (D’) and PHACTR4 (D’’). Inducible T-REx 293 cells lines were stimulated overnight with doxycycline. Pull-downs performed employing normal IgG or antibody against HA tag were compared using Western blot, N=3. **E.** Confocal imaging of the endogenous RAI14 (B) and EPHA2 (B’) proteins in A375 melanoma cells. Knock-out cell lines derived by CRISPR/Cas9 served as a negative control (Suppl. Fig. 5B). F-actin is stained by phalloidin conjugated with Alexa488 and DNA by DAPI. Scale bar represents 10 μm. **F.** Quantification of the Western blot results comparison of MLC2 phosphorylation events in A375 parental and *RAI14*, *EPHA2* and *PHACTR4* KOs (Western blot is presented in the Suppl. Fig. 5C). **G.** Measurements of cellular circularity. Fixed and crystal violet stained cells were measured and then results were analyzed using SuperPlotsOfData, N=4. At least 470 cells were measured for each cell line in single repetition. Mean results of each repetition are plotted and tested by one-way Anova, N=4; *p* values are indicated in the graph. Examples of A375, *RAI14* and *PHACTR4* KO cells are presented (G’).

WNT5A signaling is crucial for 2D migration of A375 melanoma cells, which can be analyzed in a wound healing assay (Fig. 5B and 5B’). Strikingly, A375 cells deficient in endogenous RAI14, EPHA2 or PHACTR4 showed an impaired migration in this assay and phenocopied loss of *WNT5A* (Fig. 5B-B’). WNT5A controls A375 cell migration via DVL-RhoA-ROCK-myosin light chain kinase (MLCK)-myosin light chain (MLC) cascade (schematized in Fig. 5C). Inhibition at the level of (i) WNT5A by the porcupine inhibitor (PORCNi); LGK-974, (ii); DVL by casein kinase 1 (CK1) δ/ε inhibitor (CK1i) PF-670462; (iii) ROCK by ROCK kinase inhibitor Y-27632 or (iv) inhibitor of nonmuscle myosin II (MYL2, also MLC2) Blebbistatin efficiently blocks migration in WT cells but do not further reduce migration of *RAI14*, *EPHA2* and *PHACTR4* KO lines (Fig. 5C-C’). This suggests that the migration in these KO cells is blocked by the same mechanism.

All the KO lines responded well to WNT5A as visualized by phosphorylation of DVL (see Fig. 5A). Interestingly, the steady state level of DVL2 phosphorylation was increased in all tested *RAI14* and *EPHA2* KO clones, but not in PHACTR4 deficient cells (Suppl fig 5A). We hypothesized that this may be a compensatory mechanism due to impaired signal transduction to the downstream effectors. To test this possibility, we have looked in more detail into possible interaction of RAI14, EPHA2 and PHACTR4 with downstream effectors, namely ROCK and MLC. In the co-IP assay we could show that inducibly expressed HA-tagged RAI14, EPHA2 and PHACTR4 interact with endogenous ROCK1 and pMLC (Fig. 5D-D’’). Endogenous RAI14 co-localized in A375 cells with the F-actin fibers localized to the trailing edge (Fig. 5E), whereas EPHA2 could be detected in some regions of the plasma membrane and in filopodia (Fig. 5E’) (for the validation of antibodies on KO cells see Suppl. Fig. 5B).

Phosphorylation of MLC2 by MLCK activated by ROCK affects actin network contractility^34–36^. Indeed, MLC2 phosphorylation in A375 cells could be induced by WNT5A in parental but not in the *RAI14* KO and *EPHA2* KO cells; *PHACTR4* KO showed much weaker phenotype and one KO clone responded similarly to WT (Fig. 5F and 5F’).

We have observed that some of our KO cell lines appear to be spreading less over the surface compared to parental cell line in 2D culture. Thus, we quantified the overall shape of the A375 parental cells in 2D culture and compared it with KO cells using image analysis. Indeed, lines lacking RAI14 and EPHA2 showed increased circularity compared to parental A375 cell line (Fig. 5G and 5G’), what we connected with the actomyosin network contractility (Fig. 5F and 5F’).

We conclude that RAI14, EPHA2 and PHACTR4 are essential for WNT5A-induced migration in melanoma cells, where they mediate the signal transduction between receptor complexes and actomyosin. In combination with proteomic and zebrafish data we propose RAI14, EPHA2 and PHACTR4 as novel components of Wnt/PCP pathway in vertebrates.

## DISCUSSION

The primary goal of our study was to identify novel essential components of Wnt/PCP pathway in vertebrates. In order to achieve this goal, we have mapped the interactome of five key Wnt/PCP components: ROR1, ROR2, VANGL2, PRICKLE1 and DVL3 by proximity-based labelling. By combining the interactomes of these five proteins, we could narrow down a list of candidates for functional testing. This approach allowed us to identify novel Wnt/PCP pathway proteins that were missed in the earlier proteomic studies that either used affinity purification or focused only on one bait^37–39^.

Proximity-based approaches such as BioID come with their own pros and cons. One of the disadvantages is the labeling and detection of proteins that participate in the protein biosynthesis and not on the function of the mature protein. We believe that in our study this is the case of #C2 preys, which interacted only with VANGL2. VANGL2 is a four transmembrane, tetraspanin-like, protein and its #C2 interactors localized almost exclusively to the Golgi-ER membrane network, where maturation of VANGL2 takes place. #C2 proteins thus represent candidate proteins that participate on the synthesis and post-translational modification of VANGL2, as demonstrated for example by SEC24B, protein with the previously characterized key role in VANGL2 sorting^40–42^ that falls in our dataset into cluster #C2.

On the other side, the advantage of BioID is the capacity to catch also transient interactions, and availability of the reference tools such as Human Cell Map^20^, which allows rapid and unbiased insight into localization of individual preys in the subcellular compartments. Surprisingly, even in T-REx 293 cells, which do not form typical epithelium, preys from #C3 and #C4 clusters associated almost exclusively with the compartment annotated as cell junction. This is despite the fact that TREx 293 cells do not form epithelial sheet with a globally polarized character like the typical models used to study PCP, and suggests that the proteins mediating Wnt/PCP signaling in this non-epithelial model still connect with the cell adhesion complexes and perhaps also signal in the contact-dependent manner. While cell junctions have been firmly established as the main PCP domains based on crucial studies on Drosophila wing epithelium, the external regulation of PCP at the cell junction by Wnt ligands have been at least in this particular model a matter of controversy^43^. Our proteomic data that place Wnt-binding ROR1/2 co-receptors as well as their signaling complexes to cell junction, support the possibility that it is indeed the place where non-canonical Wnt ligands signal. This is in line with the proven role of Wnt ligands in other processes – such as the contact inhibition of locomotion, which involves transient cell-cell contacts and depends critically on Wnt/PCP^44,45^ or the evidence for the contact-dependent transmission of Wnt ligands by ROR1/2 co-receptors via cytonemes^46–48^.

Detailed look into the identified preys with the known function and cell junction localization revealed multiple proteins associated with the apicobasal (AB) epithelial polarity. More specifically, with the proteins MARK2 (a PAR1 homolog), SCRIB, DLG1 and DLG5 that form its basolateral component^49^. One of the established properties connecting proteins critical for AB and PCP polarity pathways is the presence of PDZ domains, which seem to represent a common interaction interface in polarity complexes^50^. Some of the crucial polarity proteins (i. e. VANGL and SCRIB) can serve as scaffolds^51,52^, which help them to organize local signaling centers. Identification or confirmation of AB proteins as PCP protein interactors, together with identification of additional PDZ-containing proteins as preys (i. e. AHNAK2, PDLIM1 and 5, PTPN13), further strengthens the notion that Wnt/PCP represents an important part of the interconnected web of PDZ proteins regulating the global morphology of cells within the tissue.

The functional validation of the proteins that interacted with Wnt/PCP proteins in our BioID assay, but were not previously associated with the Wnt/PCP pathway, identified three proteins - RAI14, EPHA2 and PHACTR4 - as essential components of the Wnt/PCP pathway. Disruption of genes encoding these proteins resulted in the convergent extension phenotypes in zebrafish, disruption of PCP-dependent polarity in the lateral organ of zebrafish, and loss of migratory properties in melanoma cells. Interestingly, absence of none of these proteins affected WNT5A signal perception, as far as we can see on the WNT5A-induced DVL phosphorylation assay, but blocked signal transduction to downstream effectors or effectors themselves. Not surprisingly the functional consequences of the loss of individual proteins were not identical – only in the absence of RAI14 and EPHA2, which both have been previously associated with membrane and cytoskeleton, and show more similar interactome (Fig. 4D), WNT5A-induced phosphorylation of MLC was affected; this phenotype co-appeared with the more round morphology of RAI14 and EPHA2 KO cells.

RAI14 was described as a protein associated with F-actin structures^25^ that can regulate dynamics of the actin cytoskeleton^53^. It has been mechanistically linked with many cellular processes such as myosin II regulation, dendritic spine formation, macropinocytosis and regulation of cell motility, epithelial to mesenchymal transition (EMT) and proliferation^54–57^. We have observed polarized localization of the RAI14 in melanoma cells in the structures that probably correspond to WRAMP^58^, characterized by high myosin activity^59^. Recent study by Jeong and colleagues^60^ suggests that RAI14 can mediate mechanotransduction and link mechanical forces with the activation of Hippo (YAP/TAZ) signaling. This is in line with our miniTurboID-based RAI14 interactome where we see enrichment in the Hippo pathway components. Given the fact that RAI14 can also shape the local membrane domains^28^, RAI14 is a very interesting candidate for a protein that links Hippo, membrane changes, mechanotransduction and Wnt/PCP signaling. All these processes must integrate at the contact sites between adjacent cells where the mechanical forces are being transmitted between the cells and that need to be continually rebuilt to accommodate their mechanical properties to the ever-changing tissue geometry^61^.

In contrast to RAI14, EPHA2 is a relatively widely studied protein, primarily as a receptor in the ephrin signaling pathway. EPHA2 has been reported to be involved in several processes associated with the noncanonical Wnt pathway. First, it can bind WNT1 and interact with DVL2 and AXIN1^62^, two elements of the Wnt signalosome. It is not known if EPHA2 can interact with other Wnt ligands, but given the fact that it contains CRD domain, it definitely keeps this possibility open. Second, EPHA2 is localized in adherens junctions in an E-Cadherin-dependent manner^30^. Third, it has been connected to the regulation of cell migration and increased invasion in melanoma and prostate cells^63,64^ and mammary gland branching morphogenesis^65^. EPHA2 can promote amoeboid migration mode and activate Rho family GTPases, in particular RhoA and RhoG^64–66^. Based on these facts and our data, transmembrane EPHA2 is the ideal candidate that connects non-canonical Wnt/ROR/DVL receptor complexes with the cell junctions and downstream regulation of the small GTPases.

PHACTR4, the last from the novel Wnt/PCP effectors, is an understudied protein. It is known to interact with β-actin and Protein Phosphatase 1 (PP1)^29^. Besides that, PHACTR4 acts as tumor suppressor by regulating cell proliferation and *PHACTR4* gene is commonly deleted in multiple types of cancer^67^. Interestingly, the mouse strain called the *humpty dumpty* (*humdy*), which displays neural tube and optic fissure closure defects, carries Arg to Pro missense mutation at amino acid position 650 in *Phactr4*^68^. Open neural tube is a typical Wnt/PCP phenotype^1^. PHACTR4 can be asymmetrically localized at the leading edge of the cell and regulates directional migration by actin cytoskeleton modulation via the Rho-Cofilin pathway^70,71^, potentially placing PHACTR4 between plasma membrane receptor complexes and downstream effectors of the Wnt/PCP pathway. In contrast to *EPHA2* and *RAI14* KO melanoma cells, loss of PHACTR4 does not prevent WNT5A-induced phosphorylation of MLC2, which suggest that mechanistically PHACTR4 acts differently; a notion that is also supported by the analysis of its interactome that overlaps least from the studied PCP proteins. The precise mechanism of action of PHACTR4 in the Wnt/PCP will thus require further studies.

Our study provides a comprehensive proteomic dataset that identifies interactors of five proteins of the Wnt/PCP pathway: ROR1, ROR2, VANGL2, PRICKLE1 and DVL3. We focused and functionally screened in zebrafish a group of preys, which resulted in the identification of three novel protein components of vertebrate Wnt/PCP pathway. Our study is not exhaustive but provides valuable starting point for future analysis focused on the bait-specific functions, biosynthesis or the biology associated with the phase-separated cytoplasmic proteins DVL3 and/or PRICKLE.

## ACKNOWLEDGEMENTS

This project has been supported by the Czech Science Foundation (GA22-25365S) and the project National Institute for Cancer Research (Programme EXCELES, ID Project no. LX22NPO5102) - Funded by the European Union - Next Generation EU. CIISB, Instruct-CZ Centre of Instruct-ERIC EU consortium, funded by MEYS CR infrastructure project LM2023042, is gratefully acknowledged for the financial support of the measurements at the CEITEC Proteomics Core Facility. Computational resources were provided by the e-INFRA CZ project (ID:90254), supported by MEYS CR.

## MATERIALS AND METHODS

### Cell culture

T-REx-293 (R71007, Thermo Fisher Scientific), A375 and their derivate were cultured in the Dulbecco’s modified Eagle’s medium (DMEM, 41966-029, Gibco, Life Technologies) with the 10% fetal bovine serum (FBS, 10270-106, Gibco, Life Technologies), 2 mM L-glutamine (25030024, Life Technologies), 1% penicillin-streptomycin (XC-A4122/100, Biosera) under 5% (vol/vol) CO2 controlled atmosphere at 37°C. Regular screenings for mycoplasma contamination were conducted.

### Cells shape analysis

T-REx-293 WT or CRISPR KO cells were seeded in 24-well plates in a low density and 0,1% FBS DMEM, and let to adhere to the surface overnight. Then, the cells were washed with PBS, fixed using ice-cold methanol and stained with 0,5% crystal violet. The fixed samples were then let to dry at room temperature for 24 h. Violet-stained cell samples were then imaged using Leica DMI6000B inverted microscope in bright field mode, equipped with ORCA-Flash4.0 V3 Digital CMOS camera C13440-20CU (Hamamatsu), using 5x/0.15 HC PL FLUOTAR objective (Leica).

The acquired images were then batch-segmented using pre-trained ilastik (ver. 1.3.3.post3) pixel classification model^85^ and shape of the resulting cell masks analyzed in FIJI software (ver. 1.53f) using “Analyse particles” function^86^. Initial results were filtered for the “area” parameter value between 500 and 1600 pixels to discriminate staining artifacts and cell aggregates.

### Treatments

To inhibit the activity of endogenous Wnt ligands, cells were exposed to LGK-974 (1241454, PeproTech) at a concentration of 1μM. The noncanonical Wnt pathway was induced by 100 ng/ml recombinant human WNT5A (645-WN, R&D Systems) as described in the figure legends, for 3-hour incubation period or by WNT5A conditioned medium (CM) (control or WNT5A), produced by rat L-fibroblasts (CRL-2648, CRL-2647, CRL-2814; ATCC), at a ratio of 2:1 = CM : standard culture medium for 6 hours. Doxycycline (HY-N0565B, MedChem Express) or tetracycline (60-54-8, Santa Cruz Biotechnology) in dose 1 μg/ml were used overnight to the drive expression of POIs using TetON system. Following inhibitors have been used: 10μM Y-27632 (ROCKi, HY-10071, MedChem Express), 1μM Go6983 (PKCi, HY-13689 MedChem Express), 5μM Blebbistatin (Myosin II inhibitor, HY-13813 MedChem Express), 10μM EHop-016 (RACi, MedChem Express) and 3μM PF-670642 (CK1δ/ε inhibitor, HY-15490 MedChem Express).

### Plasmids and cloning

Inducible expression of POIs in T-REx-293 was achieved by using pcDNA4-TO backbone. N-terminal tagging with BirA* was possible thanks to amplification of Myc-BirA* from pcDNA3.1 mycBioID 35700 (a gift from Kyle Roux; Addgene # 35700) and fusion with sequences encoding POIs. Similarly, BirA*-HA was introduced to the C-terminal part of POIs using sequences from pcDNA3.1 MCS-BirA(R118G)-HA (Addgene # 36047). All BioID constructs were cloned into pcDNA4-TO backbone. RAI14^26^, EPHA2^88^ and PHCATR4 (RC206425, Origene) plasmids were used for fusion of cDNAs encoding POIs with V5-miniTurboID enzyme or with HA tag and cloning into pcDNA4-TO backbone. *In-Fusion Cloning* (Takara Bio) method was used for cloning. Other plasmids used: pcDNA3 Ror2 Flag^89^ and pcDNA3-Ror2-HA^90^.

### Transfection

Cells were transfected using jetOPTIMUS (Polypus, 101000025) reagent accordingly to the manufacturer’s instructions.

### T-REX 293 stable cell lines preparation

T-REx 293 stable cells lines were obtained by antibiotic selection - 100 µg/m Zeocin (ant-zn-1p, Invivogen) for pcDNA4-TO plasmid and limiting dilution method for clonal selection.

### CRISPR/Cas9 in cell culture

For generation of the Knock Out (KO) cell lines, plasmids encoding guide RNAs targeting *RAI14*, *EPHA2*, *PHACTR4* and *ROR2* genes were used (*RAI14* gRNA: GAATGATGACCGGCTACTGC, *EPHA2* gRNA: CTACTATGCCGAGTCGGACC, *PHACTR4* gRNA: TTGGACTGGACTAGATGATC, *WNT5A*: AGTATCAATTCCGACATCGA). gRNA sequences were cloned into plasmids pSpCas9(BB)-2A-BFP (Addgene plasmid, 64323). A375 cells were cultured according to manufacturer’s instructions and transfected by jetOPTIMUS® DNA Transfection Reagent (Polypus, 101000006) with plasmids encoding guide RNAs and the PiggyBac system (hygromycin resistance cassette and transponase). Stable cell lines were established by hygromycin B (sc-29067; Santa Cruz Biotechnology) at a concentration of 50 ug/ml for 10 days. Single cell colonies were then picked and screened by PCR or Western blotting (WB). For PCR, genomic DNA was isolated by using DirectPCR lysis reagent (301-C; Viagen Biotech), and then the fragment of genomic DNA was amplified by PCR using DreamTaq DNA polymerase (EP0705, Thermo Fisher Scientific). All the clones were sequenced using the Illumina platform and compared with the reference sequence^91^. Sequencing results are presented in Supplementary Table 7.

### Western blot

Western blot analysis was carried out following established protocol using samples with equal protein quantities determined through the DC Protein Assay (5000111, Bio-Rad) or direct lysis in the sample buffer (comprising 2% SDS, 10% glycerol, 5% β-mercaptoethanol, 0.002% bromophenol blue, and 0.06 M Tris HCl, pH 6.8). Briefly, following electrophoretic separation in an SDS-PAGE gel, proteins were transferred onto an Immobilon-P PVDF Membrane (IPVH00010, Millipore), nonspecific interactions were blocked in 5% non-fat milk or 3% BSA in washing buffer. Detection of proteins were subsequently performed utilizing primary antibodies along with their corresponding HRP-conjugated secondary antibodies. The Fusion SL imaging system (Vilber) and Immobilon Western Chemiluminescent HRP Substrate (Merck, WBKLS0500) were used for detection and visualization. List of antibodies is in the Supplementary Table 11. Molecular mass markers are indicated in the figures.

### Immunofluorescence and confocal microscopy

Cells cultured on glass coverslips were fixed by 4% paraformaldehyde (PFA) solution in PBS. Fixed cells were permeabilized by 0.1% Triton X-100 in PBS and blocked in 1% s bovine serum albumin (BSA) in PBS. Afterward, the specimens were subjected to an overnight incubation at 4°C with primary antibodies in a 1% BSA solution in PBS. After that, three washes with PBS were performed and corresponding Alexa Fluor secondary antibodies (Invitrogen) were incubated with samples for 1 hr at RT, along with DAPI for nuclei counterstaining. After PBS washes, the samples were mounted in the DAKO mounting medium (S3023, DAKO). Images were taken on the confocal laser scanning microscope Leica SP8. List of antibodies is in the Supplementary Table 11. Scale bars are indicated in figures.

### Wound healing assay

A375 and their derivate at 90% confluence were cultured overnight in DMEM supplemented with 0.1% FBS to induce proliferation arrest. Subsequently, the cells were subjected to mechanical wounding using a pipette tip, followed by a PBS wash. After 48 h of culture in 0.1% FBS DMEM, cells were fixed ice-cold methanol and stained with 0.5% crystal violet, washed and dried. The wound gap was documented using a Leica DMI6000 B inverted microscope. The percentage of cell-free surface area was quantified using the FIJI software^86^.

### Zebrafish crispants production

Oligonucleotide sequences for gRNA production were selected using the web tool https://chopchop.cbu.uib.no. A protocol published by Shah et al.^72^ was utilized for gRNA production. A mixture consisting of rCas9NLS protein (0.66 mg/ml, Sigma-Aldrich CAS9PL), Phenol red (0.05%, Sigma-Aldrich P0290), 120 mM KCl, and gRNA (200 ng/μl) was injected in a volume of 1nl into the cells of one-cell stage embryos of the AB zebrafish strain. In control injections, an equivalent concentration of BSA was used in place of rCAS9 in an otherwise identical mixture. After the injection embryos were kept in E3 medium at 28,5 °C. List of the gRNA sequences is available in the Supplementary Table 12.

### MyoD1 staining

At 14.5 hpf, dechorionated embryos were fixed using 4% PFA for one hour, followed by three 20-minute washes in PBST (1% Tween in PBS). Subsequently, samples were incubated for 10 minutes in 1% Triton X-100 in PBS. After permeabilization using Triton X-100, samples were washed for 20 minutes in PBST and blocked for 2 hours in 3% BSA in PBST. Following blocking, the myoD1 antibody (Abcam ab209976, 1:200) was added and incubated overnight at 4°C on a slow shaker. In the next step, samples were washed three times for 20 minutes in PBST and incubated with a secondary antibody (α-Rabbit Alexa Fluor 488, 1:500, ThermoFisher, A-11001) for 2 hours at room temperature. After incubation, a final round of washing was performed. Finally, samples were dissected, mounted on slides using Glycerol Gelatin (Sigma-Aldrich, GG1), and imaged using a Leica SP8 confocal microscope.

### Lateral line sensory hair cells staining

5 DPF old zebrafish larvae were fixed in 4% PFA for 2 hours, followed by three 40-minute washes in PBST. Subsequently, the samples were incubated for 20 minutes in 1% Triton X-100 in PBS, followed by a 40-minute PBST wash. Thereafter, the samples were blocked in 3% BSA in PBST for 4 hours. After blocking, the samples were stained using Phalloidine Alexa Fluor 488 overnight at 4°C on a slow shaker. The next step involved washing the samples three times for 20 minutes in PBST and incubating them with DAPI (1:2000, Sigma-Aldrich D9542), followed by a final 10-minute wash. The samples were then embedded on glass-bottom dishes in 1% TopVision Agarose (ThermoFisher, R0801) and imaged using a Zeiss LSM 880 with a super-resolution Airyscan detector.

### BioID and miniTurboID

BioID assay was performed accordingly to the published protocol^18^. TetON -REx™-293 BioID cells were seeded in 10cm dishes. ROR1 and ROR2 transgenes were weakly expressed even in the absence of tetracycline, so cells were not stimulated to keep their expression on the close-to the endogenous expression level, DVL3 and VANGL2 cells were treated with 50 ng/ml of tetracycline (60-54-8, Santa Cruz Biotechnology) and 400 ng/ml was used for PRICKLE1 cells. Three hours long tetracycline treatment was followed by the overnight biotin supplementation (50 μM, Santa Cruz Biotechnology). Streptavidin Sepharose High Performance beads (17-5113-01, GE Healthcare) were used for biotinylated proteins pull-down in amount of 40 μl of slurry per sample. Labeling based on the miniTurboID was done utilizing the same protocol, but with decreased time of exposure to biotin to 6 hours and doxycycline overnight stimulation in concentration 1 μg/ml (HY-N0565B, MedChem Express). The BioID experiment was performed in 6 replicates, while miniTurboID was performed in five replicates.

### Mass spectrometry data analysis (BioID)

#### On-beads digestion

Following IP washes, bead bound protein complexes were digested directly on beads by addition of 0.75 μg of trypsin (sequencing grade, Promega) in 50mM (NH4)HCO3 buffer. Beads were gently tapped to ensure even suspension of trypsin solution and incubated at 37 °C with mild agitation for two hours. Beads were then vortexed to ensure the complex falls of the beads and the partially digested complex was transferred to the clean tubes to separate it from the beads and incubated at 37 °C for 16 hours without agitation. Peptide mixtures were then cleaned by liquid-liquid extraction (3 iterations) using water saturated ethyl acetate^73^. Cleaned solutions were evaporated completely in SpeedVac concentrator (Thermo Fisher Scientific) and extracted into LC-MS vials by 2.5% formic acid (FA) in 50% acetonitrile (ACN) and 100% ACN with addition of polyethylene glycol (20,000; final concentration 0.001%)^74^ and concentrated again in a SpeedVac concentrator (Thermo Fisher Scientific) to 15μl to get rid of the ACN prior the LC-MS analyses.

#### LC-MS/MS analysis

LC-MS/MS analyses of all peptide mixtures were done using Ultimate 3000 RSLCnano system connected to Orbitrap Fusion Lumos Tribrid mass spectrometer (Thermo Fisher Scientific). Prior to LC separation, tryptic digests were online concentrated and desalted using trapping column (100 μm × 30 mm, 3.5μm particles, X-Bridge BEH 130 C18 sorbent (Waters, Milford, MA, USA; temperature of 40 °C). After washing of trapping column with 0.1% FA, the peptides were eluted (flow rate −300 nl/min) from the trapping column onto an analytical column (Acclaim Pepmap100 C18, 3 µm particles, 75 μm × 500 mm; column compartment temperature of 40 °C, Thermo Fisher Scientific) by 100 min nonlinear gradient program (1-56% of mobile phase B; mobile phase A: 0.1% FA in water; mobile phase B: 0.1% FA in 80% ACN). Equilibration of the trapping column and the analytical column was done prior to sample injection to sample loop. The analytical column outlet was directly connected to the Digital PicoView 550 (New Objective) ion source with sheath gas option and SilicaTip emitter (New Objective; FS360-20-15-N-20-C12) utilization. ABIRD (Active Background Ion Reduction Device, ESI Source Solutions) was installed.

MS data were acquired in a data-dependent strategy with defined number of scans based on precursor abundance with survey scan (m/z 350-2000 m/z). The resolution of the survey scan was 120000 (at m/z 200) with a target value of 4×105 ions and maximum injection time of 100 ms. HCD MS/MS (30% relative fragmentation energy, normal mass range) spectra were acquired with a target value of 5.0x104 and resolution of 15 000 (at m/z 200). The maximum injection time for MS/MS was 22 ms. Dynamic exclusion was enabled for 30 s after one MS/MS spectra acquisition. The isolation window for MS/MS fragmentation was set to 1.2 m/z.

#### DDA data processing

The analysis of the mass spectrometric RAW data files was carried out using the MaxQuant software (version 1.6.0.16) using default settings unless otherwise noted. MS/MS ion searches were done against modified cRAP database (based on http://www.thegpm.org/crap) containing protein contaminants like keratin, trypsin etc., and UniProtKB protein database for Homo sapiens (https://ftp.uniprot.org/pub/databases/uniprot/current_release/knowledgebase/reference_prot eomes/Eukaryota/UP000005640/UP000005640_9606.fasta.gz; version 2018/05, number of protein sequences: 20,996). Oxidation of methionine and proline, deamidation (N, Q) and acetylation (protein N-terminus) as optional modification, and trypsin/P enzyme with 2 allowed missed cleavages were set. Peptides and proteins with FDR threshold <0.01 and proteins having at least one unique or razor peptide were considered only. Match between runs was set among all analysed samples. Protein abundance was assessed using protein intensities calculated by MaxQuant. Mass spectrometry proteomics data were deposited to the ProteomeXchange Consortium via PRIDE^75^ partner repository under dataset identifier PXD048685.

### Mass spectrometry data analysis (miniTurboID)

#### On-beads digestion

Bead bound protein complexes were processed and digested according to protocol^76^. Resulting peptides were extracted into LC-MS vials by 2.5% formic acid (FA) in 50% acetonitrile (ACN) and 100% ACN with addition of polyethylene glycol (20,000; final concentration 0.001%)^74^ and concentrated in a SpeedVac concentrator (Thermo Fisher Scientific).

#### LC-MS/MS analysis

LC-MS/MS analyses were done using RSLCnano system connected to Orbitrap Exploris 480 spectrometer (Thermo Fisher Scientific) with EASY Spray ion source (Thermo Fisher Scientific) installed. Prior to LC separation, tryptic digests were online concentrated and desalted using trapping column (300 μm × 5 mm, μPrecolumn, 5μm particles, Acclaim PepMap100 C18, Thermo Fisher Scientific). After washing of trapping column with 0.1% FA, the peptides were eluted (flow 300 nl/min) from the trapping column onto analytical column (Aurora C18, 75μm ID, 250 mm long, 1.7 μm particles, heated to 50°C, Ion Opticks) by 60 min long gradient (mobile phase A: 0.1% FA in water; mobile phase B: 0.1% FA in 80% ACN).

Data were acquired in a data-independent acquisition mode (DIA). The survey scan covered m/z range of 350-1400 at resolution of 60,000 (at m/z 200) and maximum injection time of 55 ms. HCD MS/MS (27% relative fragmentation energy) were acquired in the range of m/z 200-2000 at 30,000 resolution (maximum injection time 55 ms). Overlapping windows scheme in m/z range from 400 to 800 were used as isolation window placements – see transitions_list.xlsx fie, which is available within the PRIDE repository under identifier PXD05678 as described below for more details.

#### DIA data processing

DIA data were processed in DIA-NN^77^ (version 1.8.1) in library free mode against modified cRAP database (based on http://www.thegpm.org/crap/; 112 sequences in total) and UniProtKB protein database for Homo sapiens (htttps://ftp.uniprot.org/pub/databases/uniprot/current_release/knowledgebase/reference_pro teomes/Eukaryota/UP000005640/UP000005640_9606.fasta.gz; version 2023/09, number of protein sequences: 20,594). No optional, carbamidomethylation as fixed modification and trypsin/P enzyme with 1 allowed missed cleavages and peptide length 7-30 were set during the library preparation. False discovery rate (FDR) control was set to 1% FDR. MS1 and MS2 accuracies as well as scan window parameters were set based on the initial test searches (median value from all samples ascertained parameter values). MBR was switched on.

Mass spectrometry proteomics data were deposited to the ProteomeXchange Consortium via PRIDE^75^ partner repository under dataset identifier PXD048678.

#### Statistical analysis of MS/MS data

For the analysis of BioID data, protein intensities, reported in proteinGroups.txt file, output of Maxquant, were further analyzed using R software, v. 4.2.2 and Differential Enrichment analysis of Proteomics data (DEP)^78^ package. Firstly, contaminants including proteins from cRAP database, keratins, and proteins labeled as Reverse or Only.Identified.By.Site were filtered out. Additionally, a combined list of proteins from 8 representative experiments employing BirA*-FLAG from the CRAPOME^18^ database, which were present in more than 5 experiments with average spectra value >20, was used as a background to exclude proteins unspecifically binding BirA* ligase itself. Remaining proteins were then subjected to further analysis. Protein intensities were log_2_ transformed and only proteins present in >=4 out of 6 replicates of at least one condition (bait) were retained. Intensities were normalized using variance stabilizing normalization (vsn) and missing values were imputed by random draws from a manually defined left-shifted Gaussian distribution originated from the actual data with shift of 1.8 and scale 0.3 in a column-wise manner. Limma test was used for differential expression analysis, and default DEP’s method (fdrtools) was used to correct adjust *p* values for multiple testing. Proteins were denoted as significantly upregulated and considered as interactors when passing the threshold of logFC > 1 and adjusted p-value <0.05.

For the analysis of miniTurboID data, protein MaxLFQ intensities reported in the DIA-NN main report file were further processed using the software container environment (https://github.com/OmicsWorkflows), version 4.7.7a. Processing workflow is available on WorkflowHub under the identifier 1083 (https://workflowhub.eu/workflows/1083) with a DOI https://doi.org/10.48546/WORKFLOWHUB.WORKFLOW.1083.1. Briefly, it covered: a) removal of low-quality precursors and contaminant protein groups, b) protein group intensities log_2_ transformation and LoessF normalization, c) imputation of missing values by random draws from a manually defined left-shifted Gaussian distribution originated from the actual data with shift of 1.8 and scale 0.3 in a column-wise manner, d) differential expression analysis using LIMMA statistical test calculated for protein groups quantified in at least 4 replicates of at least one sample type.

NMF map in Human Cell Map^20^ resource was used to assign the cellular localization for upregulated preys. UpSetR^79^ package was used to create UpSet plots. Heatmaps in 2D and Suppl. 4E were created using the MetaScape tool, <u>v3.5.20230101</u> or <u>v3.5.20240101</u>, respectively. Gene ontology was performed by the gprofiler2 tool, v. e107_eg54_p17_bf42210, using default settings.

Saint probability for bait-prey interactions was computed using the REPRINT^20^ resource, v. 2.0, using default settings if not stated otherwise: experiment type was set to Proximity Dependent Biotinylation; File Type as tab-separated matrix; none additional controls were selected. To compute the probabilistic SAINT score, SAINTExpress was used. For dotplots, only proteins with SAINT score > 0.9 in at least one bait-prey comparison were considered.

All scripts to reproduce the analyses are publicly available in the github repository https://github.com/bryjalab/2023_Wnt-PCP, with the DOI: 10.5281/zenodo.13903795.

### Large language models

Large language models (LLMs) were used exclusively for grammar corrections and troubleshooting of python scripts (ChatGPT 4 and 4o). No presented data was analyzed by LLMs.

## SUPPLEMENTARY MATERIALS

This article contains supplementary information:

### Supplementary figures

**Supplementary Figure 1.**
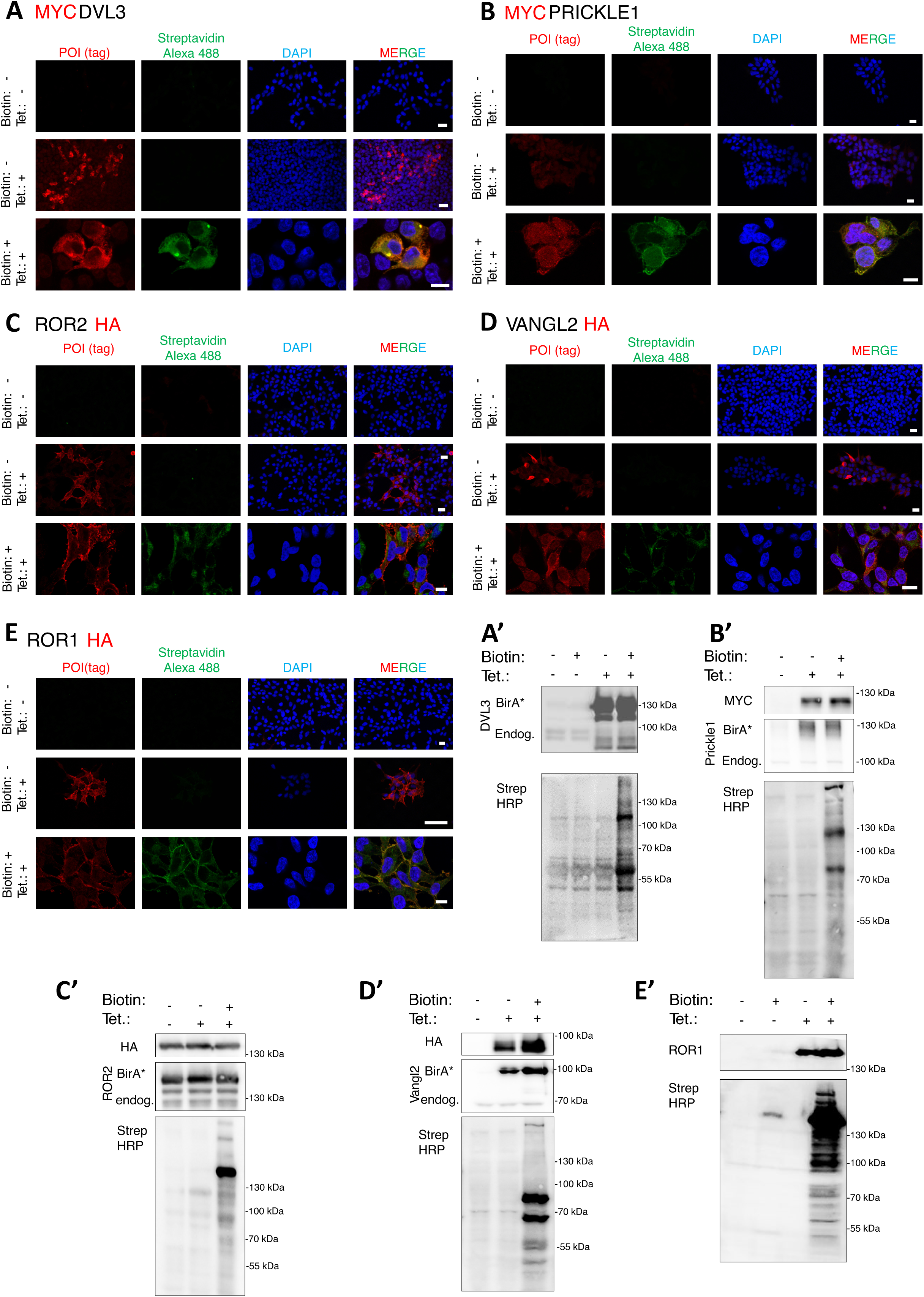
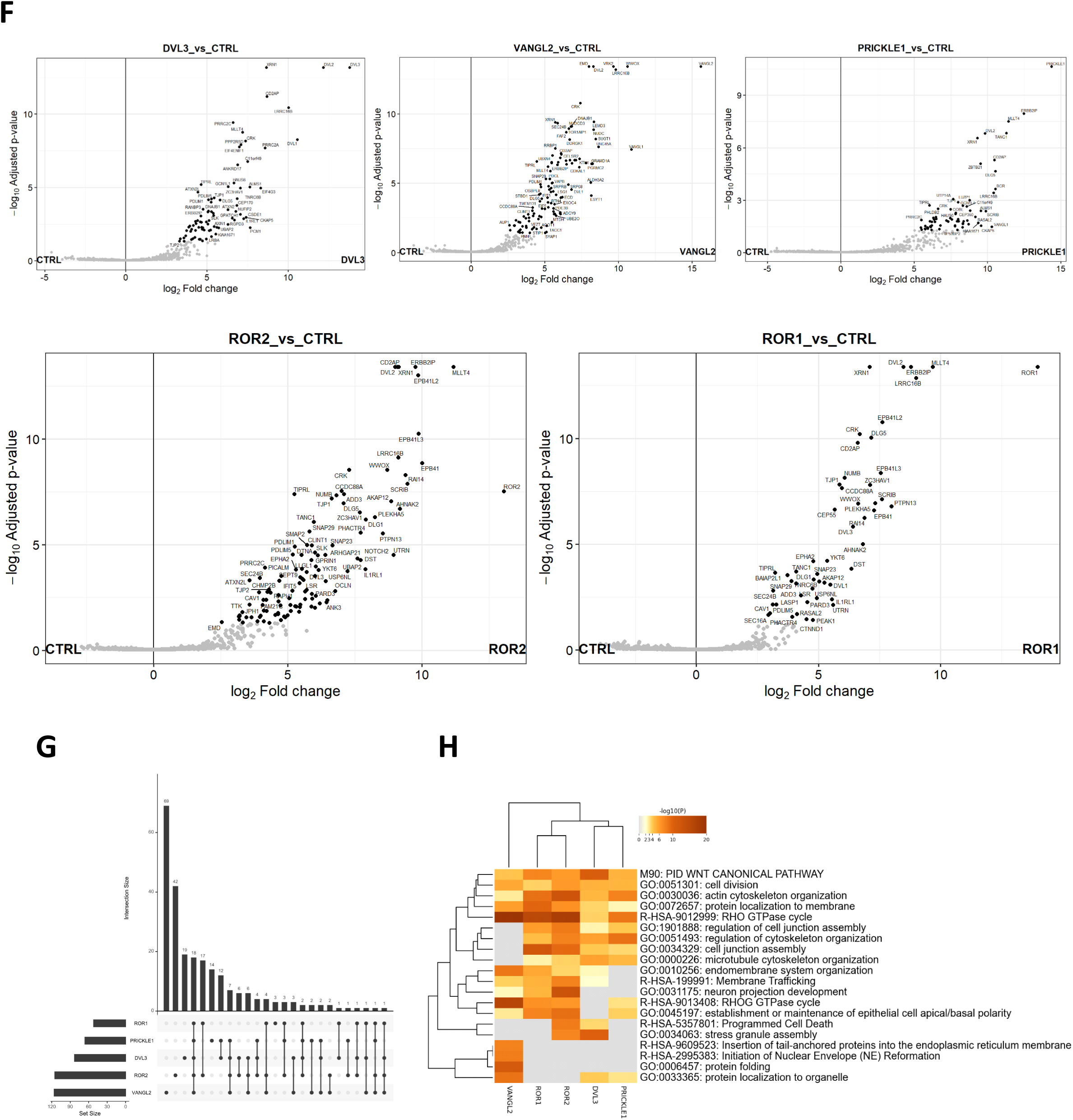
**A. - E.** Immunofluorescence analysis of baits (A MYC-DVL3; B MYC PRICKLE1, C ROR2- HA, D VANGL2-HA, E ROR1-HA) localization upon treatment with biotin and tetracycline. Streptavidin-Alexa 488 was used for biotinylation reaction products visualization. BioID cell lines were also verified by Western blot to confirm induction and BirA* reaction specificity (A’ MYC-DVL3; B’ MYC PRICKLE1, C’ ROR2-HA, D’ VANGL2-HA, E’ ROR1-HA). BirA* fusion proteins showed slower electrophoretic migration than endogenous proteins. Scale bars represent 10 μm. **F.** Volcano plots for each bait with upregulated proteins labeled. **G.** UpSet plot showing the overlap of upregulated preys in all baits. VANGL2, ROR2 and DVL3 have most unique interactomes, and there are 16 proteins which overlap in all 5 baits. **H.** Metascape analysis showing distinct gene ontology enrichment of individual bait interactors. In dendrogram, a representative term with the lowest p-value is shown, coloring corresponds to −log10(p-value), white cells indicate the lack of enrichment for particular term in given protein list.

**Supplementary Figure 2.**
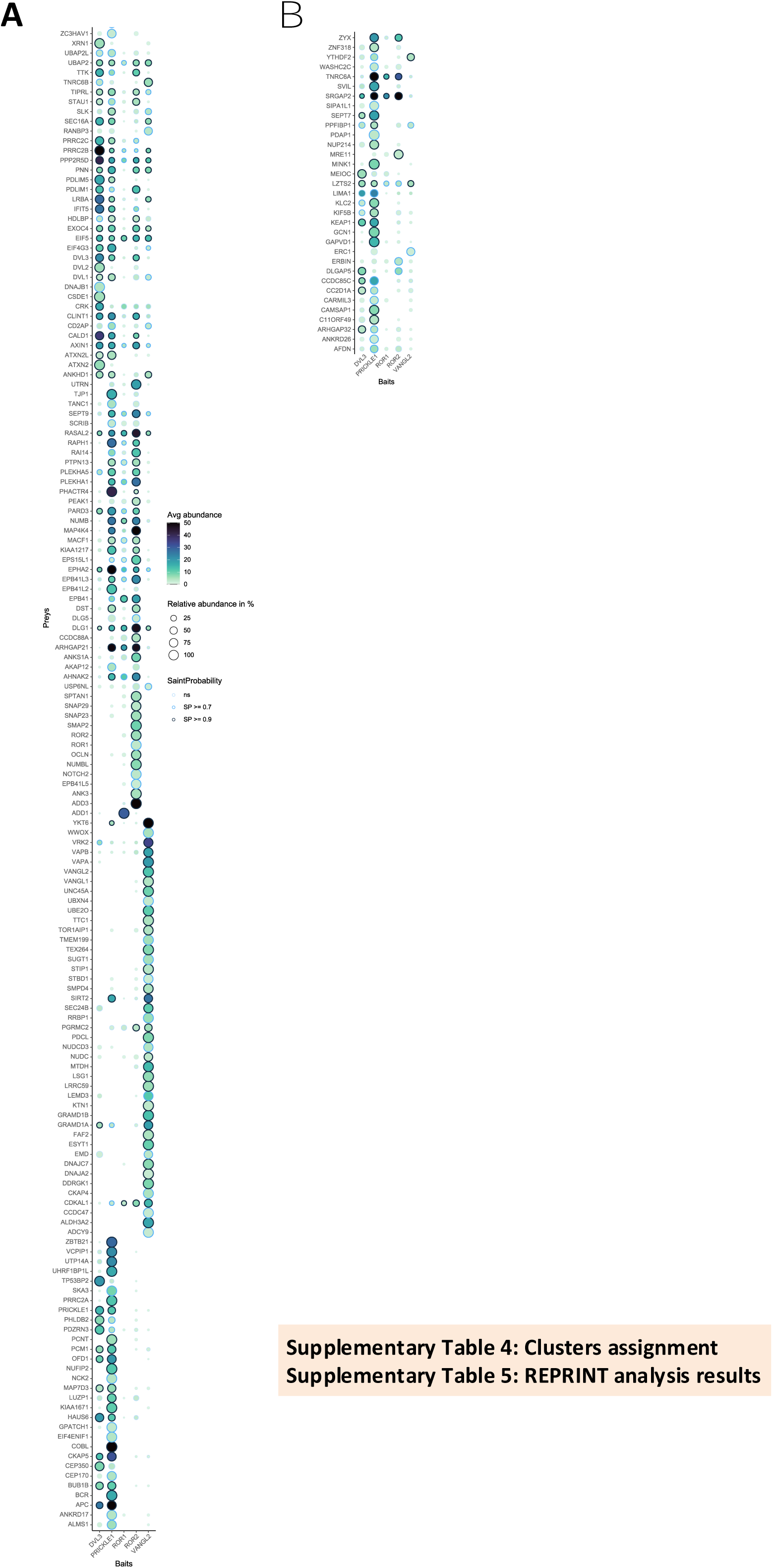
**A.** DotPlot as provided by the ProHits visualization tool, with original clustering algorithm applied, not separated by the clusters. **B.** DotPlot for not significant proteins.

**Supplementary Figure 3.**
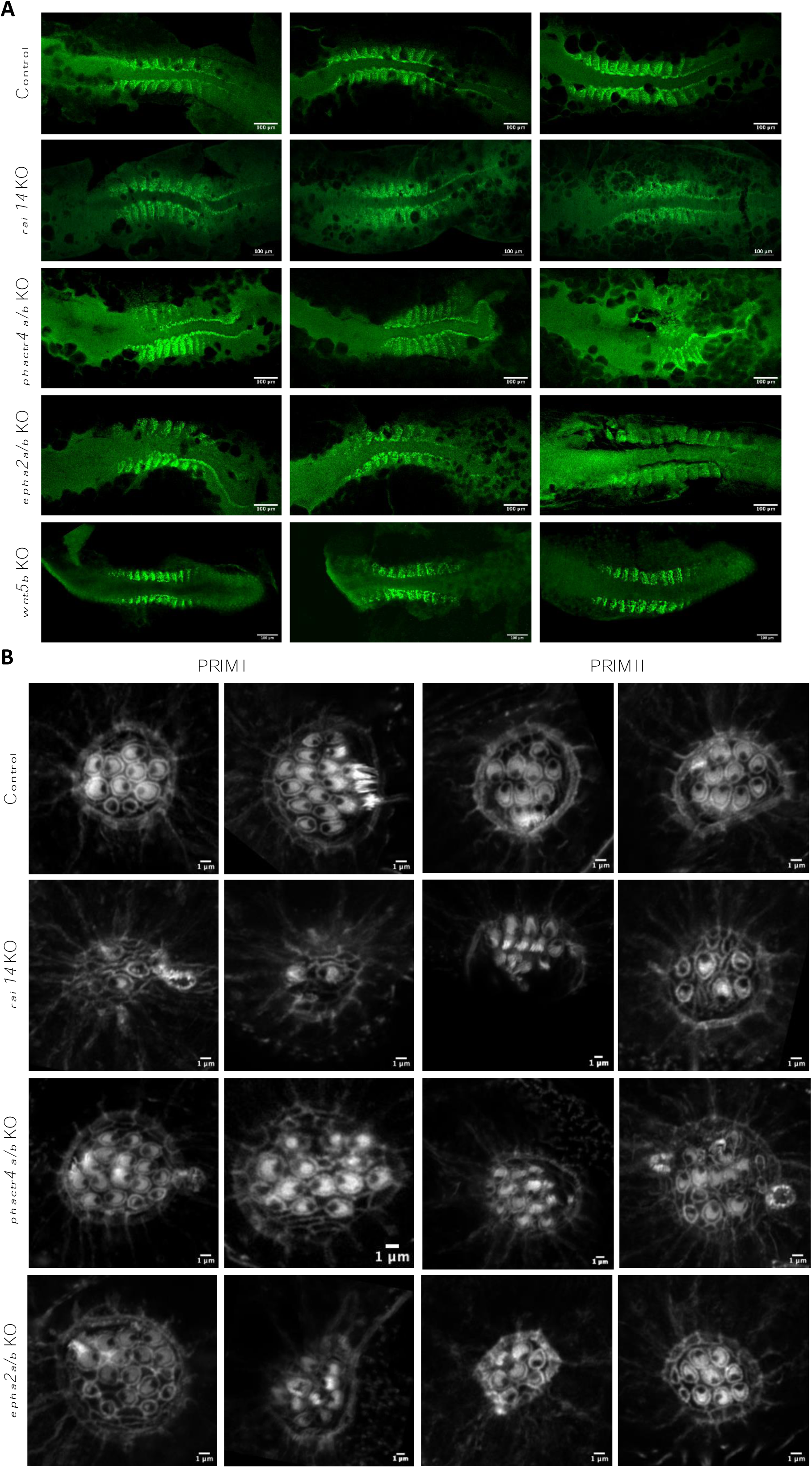
**A.** Representative images of myoD1 whole mount immunofluorescence, labeling somitogenesis in control, positive control wnt5b, rai14 KO, phactr4a/b KO and epha2a/b KO 14.5 hpf zebrafish embryos. **B.** Representative images of sensory hair cells of PRIM I and PRIM II derived neuromasts in control, rai14, phactr4a/b and epha2a/b KO embryos.

**Supplementary Figure 4.**
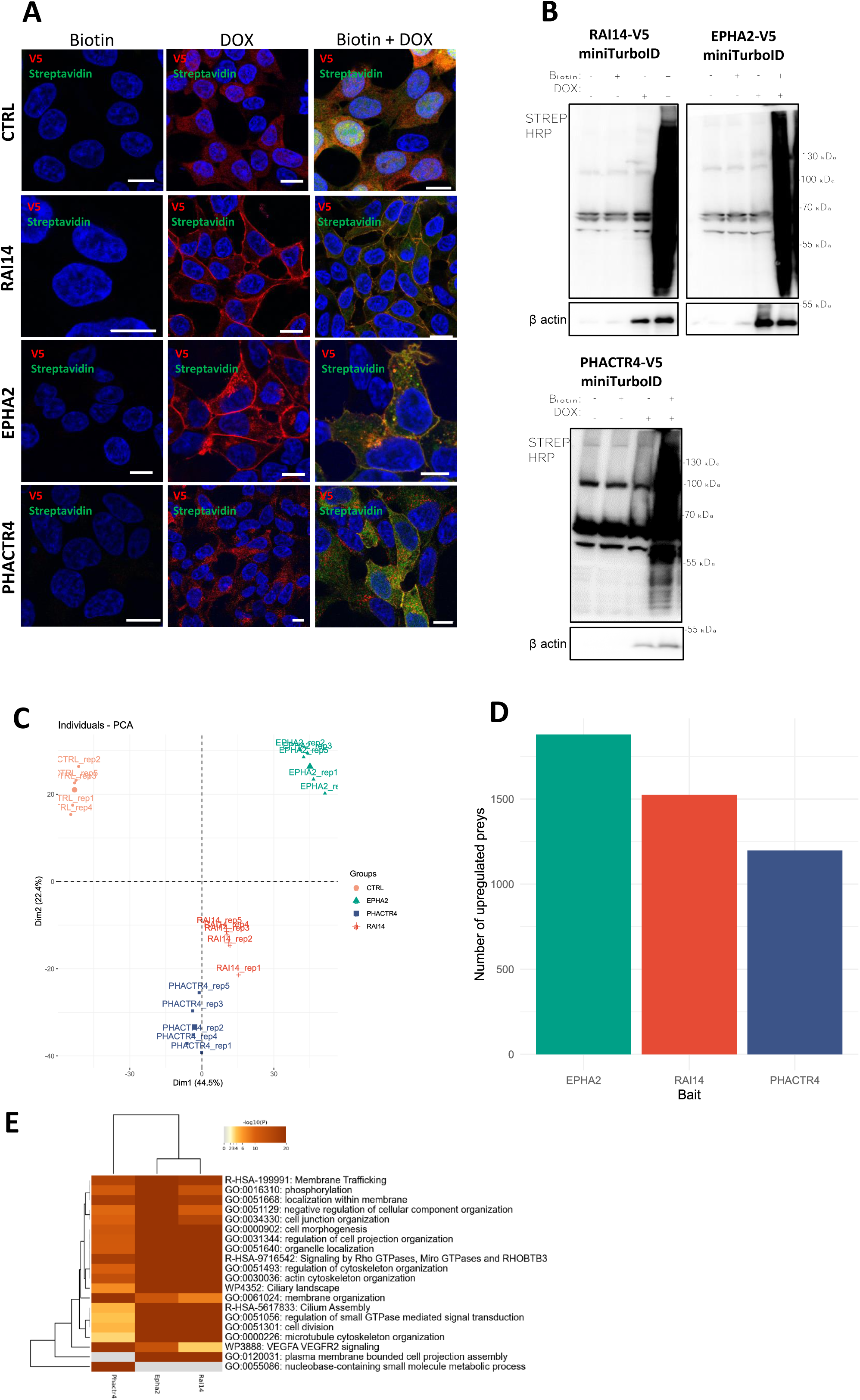
**A.** Cells inducibly overexpressing V5-miniTurbo alone or fused with cDNA encoding RAI14, EPHA2 and PHACTR4 were visualized to determine biotinylation reaction specificity and robustness. The presence of transgene is visualized by anti V5 tag antibody, labeling products by Streptavidin-Alexa488 conjugate and DNA by DAPI counterstaining. In the presence of doxycycline (DOX) treatment transgenes are expressed, but miniTurboID reaction is absent. Only co-treatment with biotin and doxycycline allows biotinylation reaction to occur. Scale bars represent 10 μm. **B.** Western blot analysis of the reaction facilitated by the POIs fused to V5-miniTurboID enzyme. Streptavidin-HRP (Strep-HRP) helped to detect biotinylation reaction products and β-actin is a loading control. Doxycycline treatment was performed overnight, and biotin supplementation lasted 6h before cell lysis. **C.** Principal Component Analysis (PCA) showing clear separation of the clusters based on the baits. **D.** Number of upregulated preys for individual baits. **E.** Gene ontology terms of RAI14, EPHA2 and PHACTR4 interactomes.

**Supplementary Figure 5.**
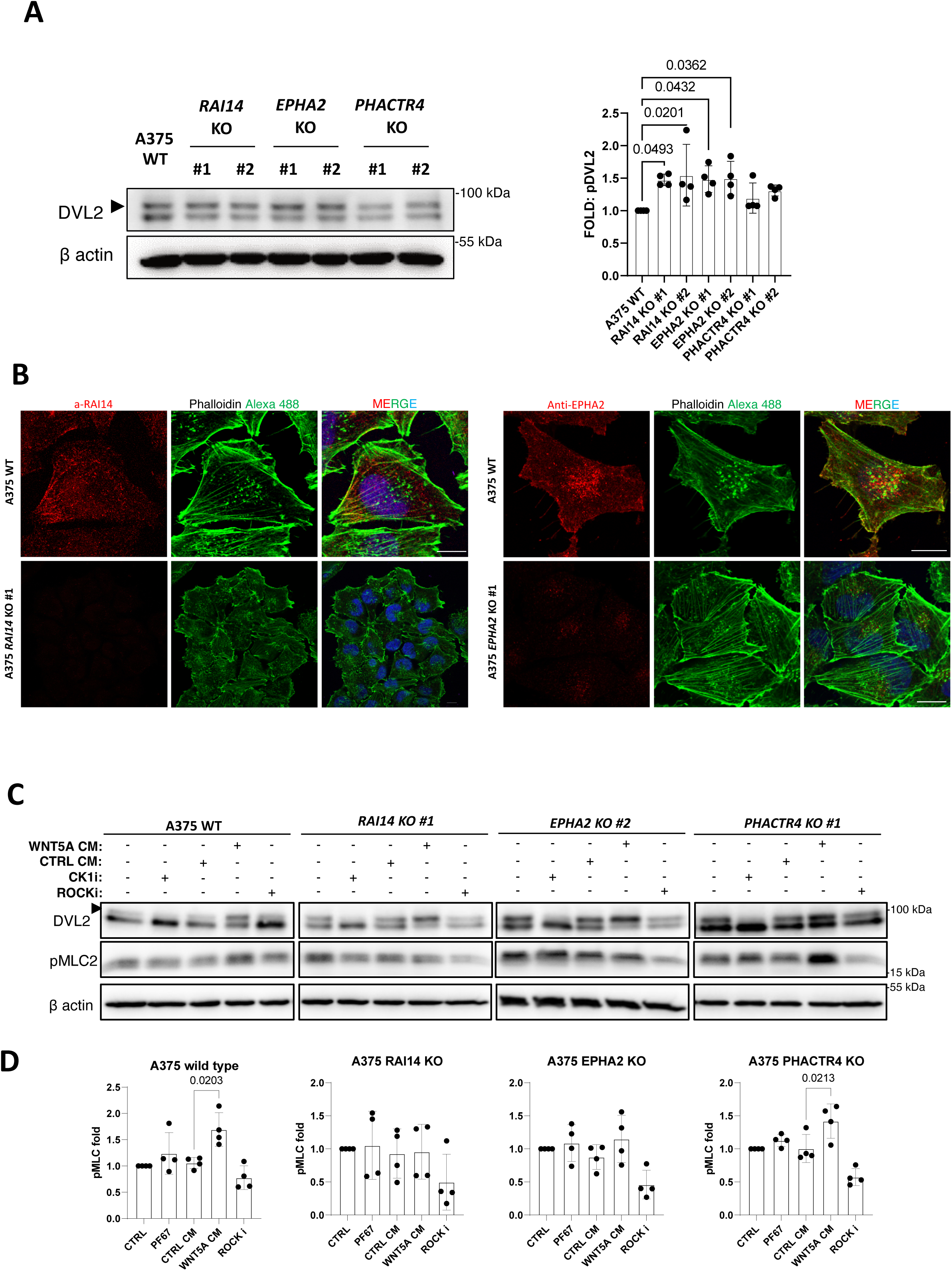
**A.** Western blot and quantification (A’) of the DVL2 phosphorylation. One-way Anova, N=4. **B.** Continuation of the a-RAI14 and a-EPHA2 antibodies validation. Confocal imaging of the endogenous RAI14 (B) and EPHA2 (B’) proteins in A375 KO melanoma cells. F-actin is stained by phalloidin conjugated with Alexa488 and DNA by DAPI. Scale bar represents 10 μm. **C.** A375 wild type and *RAI14*, *EPHA2* and *PHACTR4* KO lines were treated with WNT5A (WNT5A CM) or control conditioned media (CTRL CM) for 6 hours in 2:1 ratio (CM : 10% DMEM). Inhibitors - 10μM Y-27632 (ROCKi) or 3μM PF-670462 (CK1i) were used as negative control of DVL2 and pMLC readouts, N=4. **D.** Quantitative data showing response of tested cell lines to WNT5A CM on the pMCL level is presented (E’). More quantification data is shown in the Supplementary Figure 5A, one-way Anova, N=4.

### Supplementary tables

https://zenodo.org/records/14651166

Supplementary Table 1 Statistics results for each protein in the BioID screen

Supplementary Table 2 Significantly upregulated preys for each bait. Proteins were sorted in descending manner by log_2_FoldChange.

Supplementary Table 3 UpSet plot background table

Supplementary Table 4 Clusters assignment based on the heatmap

Supplementary Table 5 REPRINT (SaintExpress) analysis results

Supplementary Table 6 NGS results for fish

Supplementary Table 7 NGS results for the KO lines

Supplementary Table 8 Statistics results for the TurboID of 3 hits

Supplementary Table 9 Significantly upregulated preys for each bait. Proteins were sorted in descending manner by log_2_FoldChange.

Supplementary Table 10 Interaction of RAI14/EPHA2/PHACTR4 with proteins from the cluster 4

Supplementary Table 11 List of antibodies

Supplementary Table 12 List of oligonucleotides used for Zebrafish KO production

## REFERENCES

1. Wansleeben, C. & Meijlink, F. The planar cell polarity pathway in vertebrate development. Developmental Dynamics 240, 616–626 (2011).

2. Adler, P. N. Planar Signaling and Morphogenesis in Drosophila. Developmental Cell 2, 525– 535 (2002).

3. Goodrich, L. V. & Strutt, D. Principles of planar polarity in animal development. Development 138, 1877–1892 (2011).

4. Yang, Y. & Mlodzik, M. Wnt-Frizzled/Planar Cell Polarity Signaling: Cellular Orientation by Facing the Wind (Wnt). Annu. Rev. Cell Dev. Biol. 31, 623–646 (2015).

5. Lapébie, P., Borchiellini, C. & Houliston, E. Dissecting the PCP pathway: One or more pathways?: Does a separate WntLFzLRho pathway drive morphogenesis? BioEssays 33, 759–768 (2011).

6. VanderVorst, K. et al. Wnt/PCP Signaling Contribution to Carcinoma Collective Cell Migration and Metastasis. Cancer Research 79, 1719–1729 (2019).

7. Wu, J. & Mlodzik, M. The Frizzled Extracellular Domain Is a Ligand for Van Gogh/Stbm during Nonautonomous Planar Cell Polarity Signaling. Developmental Cell 15, 462–469 (2008).

8. Sebbagh, M. & Borg, J.-P. Insight into planar cell polarity. Experimental Cell Research 328, 284–295 (2014).

9. Devenport, D. The cell biology of planar cell polarity. Journal of Cell Biology 207, 171–179 (2014).

10. Jones, C. & Chen, P. Planar cell polarity signaling in vertebrates. BioEssays 29, 120–132 (2007).

11. Endo, M., Nishita, M. & Minami, Y. Analysis of Wnt/Planar Cell Polarity Pathway in Cultured Cells. in Planar Cell Polarity (ed. Turksen, K.) vol. 839 201–214 (Springer New York, New York, NY, 2012).

12. Martinez, S. et al. The PTK7 and ROR2 Protein Receptors Interact in the Vertebrate WNT/Planar Cell Polarity (PCP) Pathway. Journal of Biological Chemistry 290, 30562–30572 (2015).

13. Topczewski, J., Dale, R. M. & Sisson, B. E. Planar cell polarity signaling in craniofacial development. Organogenesis 7, 255–259 (2011).

14. Wang, M., De Marco, P., Capra, V. & Kibar, Z. Update on the Role of the Non-Canonical Wnt/Planar Cell Polarity Pathway in Neural Tube Defects. Cells 8, 1198 (2019).

15. White, J. J. et al. WNT Signaling Perturbations Underlie the Genetic Heterogeneity of Robinow Syndrome. The American Journal of Human Genetics 102, 27–43 (2018).

16. Chen, Y., Chen, Z., Tang, Y. & Xiao, Q. The involvement of noncanonical Wnt signaling in cancers. Biomedicine & Pharmacotherapy 133, 110946 (2021).

17. Roux, K. J., Kim, D. I., Raida, M. & Burke, B. A promiscuous biotin ligase fusion protein identifies proximal and interacting proteins in mammalian cells. Journal of Cell Biology 196, 801–810 (2012).

18. Mellacheruvu, D. et al. The CRAPome: a contaminant repository for affinity purification– mass spectrometry data. Nat Methods 10, 730–736 (2013).

19. Zhou, Y. et al. Metascape provides a biologist-oriented resource for the analysis of systems-level datasets. Nat Commun 10, 1523 (2019).

20. Go, C. D. et al. A proximity-dependent biotinylation map of a human cell. Nature 595, 120– 124 (2021).

21. Teo, G. et al. SAINTexpress: Improvements and additional features in Significance Analysis of INTeractome software. Journal of Proteomics 100, 37–43 (2014).

22. Navajas Acedo, J., et al. PCP and Wnt pathway components act in parallel during zebrafish mechanosensory hair cell orientation. Nat Commun 10, 3993 (2019).

23. Bai, Y. et al. Ror2 Receptor Mediates Wnt11 Ligand Signaling and Affects Convergence and Extension Movements in Zebrafish. Journal of Biological Chemistry 289, 20664–20676 (2014).

24. Ghysen, A. & Dambly-Chaudière, C. The lateral line microcosmos. Genes Dev. 21, 2118– 2130 (2007).

25. Peng, Y. et al. Ankycorbin: a novel actin cytoskeletonLassociated protein. Genes to Cells 5, 1001–1008 (2000).

26. Kutty, R. K. et al. Molecular Characterization and Developmental Expression ofNORPEG, a Novel Gene Induced by Retinoic Acid. Journal of Biological Chemistry 276, 2831–2840 (2001).

27. Kim, S. J. et al. Retinoic acid-induced protein 14 controls dendritic spine dynamics associated with depressive-like behaviors. eLife 11, e77755 (2022).

28. Wolf, D. et al. Ankyrin repeat-containing N-Ank proteins shape cellular membranes. Nat Cell Biol 21, 1191–1205 (2019).

29. Favot, L., Gillingwater, M., Scott, C. & Kemp, P. R. Overexpression of a family of RPEL proteins modifies cell shape. FEBS Letters 579, 100–104 (2005).

30. Zantek, N. D. et al. E-cadherin regulates the function of the EphA2 receptor tyrosine kinase. Cell Growth Differ 10, 629–638 (1999).

31. Larsen, A. B. et al. Activation of the EGFR Gene Target EphA2 Inhibits Epidermal Growth Factor–Induced Cancer Cell Motility. Molecular Cancer Research 5, 283–293 (2007).

32. Branon, T. C. et al. Efficient proximity labeling in living cells and organisms with TurboID. Nat Biotechnol 36, 880–887 (2018).

33. Radaszkiewicz, T. et al. RNF43 inhibits WNT5A-driven signaling and suppresses melanoma invasion and resistance to the targeted therapy. eLife 10, e65759 (2021).

34. Totsukawa, G. et al. Distinct Roles of Rock (Rho-Kinase) and Mlck in Spatial Regulation of Mlc Phosphorylation for Assembly of Stress Fibers and Focal Adhesions in 3t3 Fibroblasts. The Journal of Cell Biology 150, 797–806 (2000).

35. Ikebe, M., Hartshorne, D. J. & Elzinga, M. Phosphorylation of the 20,000-dalton light chain of smooth muscle myosin by the calcium-activated, phospholipid-dependent protein kinase. Phosphorylation sites and effects of phosphorylation. Journal of Biological Chemistry 262, 9569– 9573 (1987).

36. Riento, K. & Ridley, A. J. ROCKs: multifunctional kinases in cell behaviour. Nat Rev Mol Cell Biol 4, 446–456 (2003).

37. Salokas, K., et al. Physical and functional interactome atlas of human receptor tyrosine kinases. EMBO Reports 23, e54041 (2022).

38. Raivola, J. et al. New insights into the molecular mechanisms of ROR1, ROR2, and PTK7 signaling from the proteomics and pharmacological modulation of ROR1 interactome. Cell. Mol. Life Sci. 79, 276 (2022).

39. Daulat, A. M. et al. Mink1 Regulates β-Catenin-Independent Wnt Signaling via Prickle Phosphorylation. Molecular and Cellular Biology 32, 173–185 (2012).

40. Merte, J. et al. Sec24b selectively sorts Vangl2 to regulate planar cell polarity during neural tube closure. Nat Cell Biol 12, 41–46 (2010).

41. Feng, D. et al. Regulation of Wnt/PCP signaling through p97/VCP-KBTBD7–mediated Vangl ubiquitination and endoplasmic reticulum–associated degradation. Sci. Adv. 7, eabg2099 (2021).

42. Yang, W. et al. Wnt-induced Vangl2 phosphorylation is dose-dependently required for planar cell polarity in mammalian development. Cell Res 27, 1466–1484 (2017).

43. Ewen-Campen, B., Comyn, T., Vogt, E. & Perrimon, N. No Evidence that Wnt Ligands Are Required for Planar Cell Polarity in Drosophila. Cell Reports 32, 108121 (2020).

44. Matthews, H. K. et al. Directional migration of neural crest cells in vivo is regulated by Syndecan-4/Rac1 and non-canonical Wnt signaling/RhoA. Development 135, 1771–1780 (2008).

45. Carmona-Fontaine, C. et al. Contact inhibition of locomotion in vivo controls neural crest directional migration. Nature 456, 957–961 (2008).

46. Zhang, C., Brunt, L., Ono, Y., Rogers, S. & Scholpp, S. Cytoneme-mediated transport of active Wnt5b–Ror2 complexes in zebrafish. Nature 625, 126–133 (2024).

47. Brunt, L. et al. Vangl2 promotes the formation of long cytonemes to enable distant Wnt/β-catenin signaling. Nat Commun 12, 2058 (2021).

48. Hall, E. T. et al. Cytoneme signaling provides essential contributions to mammalian tissue patterning. Cell 187, 276–293.e23 (2024).

49. Buckley, C. E. & St Johnston, D. Apical–basal polarity and the control of epithelial form and function. Nat Rev Mol Cell Biol 23, 559–577 (2022).

50. Bilder, D. PDZ proteins and polarity: functions from the fly. Trends in Genetics 17, 511–519 (2001).

51. Bonello, T. T. & Peifer, M. Scribble: A master scaffold in polarity, adhesion, synaptogenesis, and proliferation. Journal of Cell Biology 218, 742–756 (2019).

52. Dreyer, C. A., VanderVorst, K. & Carraway, K. L. Vangl as a Master Scaffold for Wnt/Planar Cell Polarity Signaling in Development and Disease. Front. Cell Dev. Biol. 10, 887100 (2022).

53. Qian, X., Mruk, D. D. & Cheng, C. Y. Rai14 (Retinoic Acid Induced Protein 14) Is Involved in Regulating F-Actin Dynamics at the Ectoplasmic Specialization in the Rat Testis*. PLoS ONE 8, e60656 (2013).

54. Lobos Patorniti, N., et al. Rai14 is a novel interactor of Invariant chain that regulates macropinocytosis. Front. Immunol. 14, 1182180 (2023).

55. Chen, C. et al. Knockdown of RAI14 suppresses the progression of gastric cancer. OTT **Volume** 11, 6693–6703 (2018).

56. Gu, M. et al. Downregulation of RAI14 inhibits the proliferation and invasion of breast cancer cells. J. Cancer 10, 6341–6348 (2019).

57. Zhang, R. et al. Phenotypic Heterogeneity Analysis of APC-Mutant Colon Cancer by Proteomics and Phosphoproteomics Identifies RAI14 as a Key Prognostic Determinant in East Asians and Westerners. Molecular & Cellular Proteomics 22, 100532 (2023).

58. Witze, E. S., Litman, E. S., Argast, G. M., Moon, R. T. & Ahn, N. G. Wnt5a Control of Cell Polarity and Directional Movement by Polarized Redistribution of Adhesion Receptors. Science 320, 365–369 (2008).

59. Witze, E. S. et al. Wnt5a Directs Polarized Calcium Gradients by Recruiting Cortical Endoplasmic Reticulum to the Cell Trailing Edge. Developmental Cell 26, 645–657 (2013).

60. Jeong, W., Kwon, H., Park, S. K., Lee, I.-S. & Jho, E. Retinoic acid-induced protein 14 links mechanical forces to Hippo signaling. EMBO Rep 25, 4033–4061 (2024).

61. Pinheiro, D. & Bellaïche, Y. Mechanical Force-Driven Adherens Junction Remodeling and Epithelial Dynamics. Developmental Cell 47, 3–19 (2018).

62. Peng, Q. et al. EPH receptor A2 governs a feedback loop that activates Wnt/β-catenin signaling in gastric cancer. Cell Death Dis 9, 1146 (2018).

63. Taddei, M. L. et al. EphA2 Induces Metastatic Growth Regulating Amoeboid Motility and Clonogenic Potential in Prostate Carcinoma Cells. Molecular Cancer Research 9, 149–160 (2011).

64. Parri, M., Taddei, M. L., Bianchini, F., Calorini, L. & Chiarugi, P. EphA2 Reexpression Prompts Invasion of Melanoma Cells Shifting from Mesenchymal to Amoeboid-like Motility Style. Cancer Research 69, 2072–2081 (2009).

65. Vaught, D., Chen, J. & Brantley-Sieders, D. M. Regulation of Mammary Gland Branching Morphogenesis by EphA2 Receptor Tyrosine Kinase. MBoC 20, 2572–2581 (2009).

66. Hiramoto-Yamaki, N. et al. Ephexin4 and EphA2 mediate cell migration through a RhoG-dependent mechanism. Journal of Cell Biology 190, 461–477 (2010).

67. Solimini, N. L. et al. STOP gene *Phactr4* is a tumor suppressor. Proc. Natl. Acad. Sci. U.S.A. 110, (2013).

68. Kim, T.-H., Goodman, J., Anderson, K. V. & Niswander, L. Phactr4 Regulates Neural Tube and Optic Fissure Closure by Controlling PP1-, Rb-, and E2F1-Regulated Cell-Cycle Progression. Developmental Cell 13, 87–102 (2007).

69. Sun, Z. & Fässler, R. A firm grip does not always pay off: a new Phact(r) 4 integrin signaling. Genes Dev. 26, 1–5 (2012).

70. Zhang, Y., Kim, T.-H. & Niswander, L. Phactr4 regulates directional migration of enteric neural crest through PP1, integrin signaling, and cofilin activity. Genes Dev. 26, 69–81 (2012).

71. Huet, G. et al. Actin-regulated feedback loop based on Phactr4, PP1 and cofilin maintains the actin monomer pool. Journal of Cell Science 126, 497–507 (2013).

72. Shah, A. N., Davey, C. F., Whitebirch, A. C., Miller, A. C. & Moens, C. B. Rapid reverse genetic screening using CRISPR in zebrafish. Nat Methods 12, 535–540 (2015).

73. Yeung, Y.-G., Nieves, E., Angeletti, R. H. & Stanley, E. R. Removal of detergents from protein digests for mass spectrometry analysis. Analytical Biochemistry 382, 135–137 (2008).

74. Stejskal, K., Potěšil, D. & Zdráhal, Z. Suppression of Peptide Sample Losses in Autosampler Vials. J. Proteome Res. 12, 3057–3062 (2013).

75. Perez-Riverol, Y. et al. The PRIDE database resources in 2022: a hub for mass spectrometry-based proteomics evidences. Nucleic Acids Research 50, D543–D552 (2022).

76. Hollenstein, D. M., et al. Acetylation of lysines on affinity-purification matrices to reduce co-digestion of bead-bound ligands. (2023).

77. Demichev, V., Messner, C. B., Vernardis, S. I., Lilley, K. S. & Ralser, M. DIA-NN: neural networks and interference correction enable deep proteome coverage in high throughput. Nat Methods 17, 41–44 (2020).

78. Zhang, X. et al. Proteome-wide identification of ubiquitin interactions using UbIA-MS. Nat Protoc 13, 530–550 (2018).

79. Conway, J. R., Lex, A. & Gehlenborg, N. UpSetR: an R package for the visualization of intersecting sets and their properties. Bioinformatics 33, 2938–2940 (2017).

